# Highly Excretable Gold Supraclusters for Translatable *In Vivo* Raman Imaging of Tumors

**DOI:** 10.1101/2022.10.18.512314

**Authors:** Jung Ho Yu, Myeong Seon Jeong, Emma Olivia Cruz, Israt S. Alam, Spencer K. Tumbale, Aimen Zlitni, Song Yeul Lee, Yong Il Park, Katherine Ferrara, Seung-Hae Kwon, Sanjiv S. Gambhir, Jianghong Rao

## Abstract

Raman spectroscopy provides excellent specificity for *in vivo* preclinical imaging through a readout of fingerprint-like spectra. To achieve sufficient sensitivity for *in vivo* Raman imaging, metallic gold nanoparticles larger than 10 nm were employed to amplify Raman signals via surface-enhanced Raman scattering (SERS). However, the inability to excrete such large gold nanoparticles has restricted the translation of Raman imaging. Here we present Raman-active metallic gold supraclusters that are biodegradable and excretable as nanoclusters. Although the small size of the gold nanocluster building blocks compromises the electromagnetic field enhancement effect, the supraclusters exhibit bright and prominent Raman scattering comparable to that of large gold nanoparticle-based SERS nanotags due to high loading of NIR-resonant Raman dyes and much suppressed fluorescence background by metallic supraclusters. The bright Raman scattering of the supraclusters was pH-responsive, and we successfully performed *in vivo* Raman imaging of acidic tumors in mice. Furthermore, in contrast to large gold nanoparticles that remain in the liver and spleen, the supraclusters dissociated into small nanoclusters, and 73% of the administered dose to mice was excreted over 4 months. The highly excretable Raman supraclusters demonstrated here offer great potential for clinical applications of *in vivo* Raman imaging by replacing non-excretable large gold nanoparticles.

---

Raman spectroscopy identifies the unique spectral fingerprint of molecules, thus providing high specificity for biomedical imaging.^1–3^ There have been significant advancements for *in vivo* preclinical Raman imaging in various applications,^4,5^ from intraoperative imaging of microscopic tumors^6–12^ to multiplexed imaging in living subjects.^13–16^ Because the Raman scattering from molecules is generally weak, gold nanoparticles are employed to amplify the Raman signals through the localized surface plasmon-generated electromagnetic field, which is known as surface-enhanced Raman scattering (SERS).^17–19^ Since the degree of the field enhancement is proportional to the size of metallic nanoparticles, larger gold nanoparticles over 50 nm in diameter have been utilized to achieve sufficient sensitivity for *in vivo* imaging.^20^ The large gold nanoparticles exhibit minimal or negligible toxicity after systematic administration.^21^ However, they remain intact in the reticuloendothelial systems of the liver and spleen for a long period of time.^22^ Such a long-term exposure of nanoparticles could affect gene expression level, limiting their translation in humans.^23^ On the other hand, the glomerular filtration in the kidney allows clearance of nanoclusters smaller than 5 nm,^24–27^ but such small nanoclusters do not generate an electromagnetic field significant for SERS.

In this work, we have addressed this limitation by assembling renally clearable gold nanoclusters into large metallic supraclusters (Figure 1a). These assembled supraclusters generated strong Raman scattering comparable to large metallic nanostructures but are efficiently cleared from the body after biodegradation to small nanoclusters (Figure 1b). Previously, gold supraclusters were developed^28^ to improve the delivery of radiosensitizing nanoclusters into tumors^29,30^ or to enhance near-infrared (NIR) absorption for photoacoustic imaging^31,32^ and photothermal therapy^33,34^ while facilitating their excretion. On the other hand, Raman scattering supraclusters have not yet been demonstrated. Since Raman scattering is particularly sensitive to the characteristics of metallic substrates, it is critical to engineer highly excretable metallic gold supraclusters for translatable Raman imaging. The gold nanocluster building blocks are in the size range where both renal clearance and the transition from molecular state to metallic state occur, which necessitate precise synthetic control of the nanoclusters.^35^ While previously synthesized supraclusters composed of molecular gold nanoclusters showed significant improvement in excretion,^30^ little progress has been made in the design of excretable metallic supraclusters, which were intended to replace large gold nanoparticles.^36,37^

**Figure 1.**
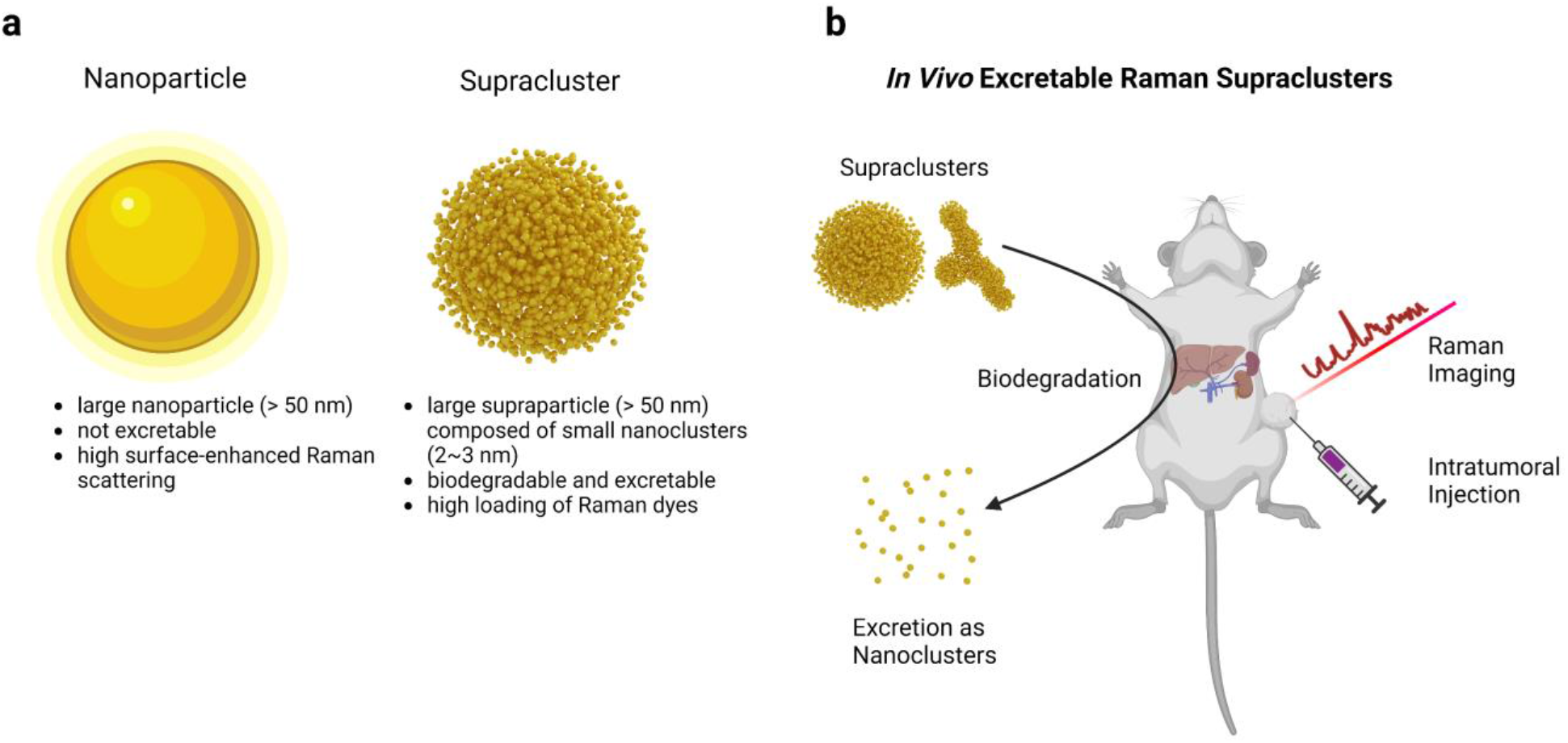
(**a**) A schematic comparison of a gold nanoparticle and a supracluster. (**b**) A scheme of spherical and branched gold supraclusters that are biodegradable and excretable as nanoclusters in the liver and spleen and perform *in vivo* Raman imaging of tumors.

Herein, we present Raman scattering gold supraclusters that provide not only high sensitivity as large gold nanoparticles-based SERS nanotags but also high excretion efficacy. The controlled synthesis of renally clearable metallic nanoclusters, their assembly into supraclusters, and the high loading of the NIR-resonant Raman dyes all contribute to Raman scattering as bright as the SERS nanotags. The supraclusters exhibit pH-dependent stability that enables *in vivo* Raman imaging of acidic tumor environments while disassembled and excreted from the liver, spleen, and kidneys. The Raman supraclusters demonstrated here thus have great potential for enabling translational *in vivo* Raman imaging.

## RESULT AND DISCUSSION

### Synthesis of renally excretable metallic gold nanoclusters

In order to produce highly excretable, Raman scattering metallic gold supraclusters, we first prepared renally clearable glutathione (ester)-capped gold nanocluster building blocks (Figure 2 and Supporting Information Figure S1 and S2). The nanoclusters were controlled to establish a size window to endow metallic properties and enable efficient clearance. The small capping ligand glutathione maintained the physiological stability of gold nanoclusters while keeping their hydrodynamic diameters small.^24^ We extended the choice of capping ligands for nanoclusters from glutathione to glutathione esters to investigate the effect of negative charges on the synthesis and properties of supraclusters. For this purpose, we synthesized 3-mercaptobenzoic acid (3-MBA)-capped gold nanoclusters with modification of the method developed by Kornberg and his coworkers.^38^ Then, the surface ligand on the nanoclusters was exchanged with glutathione monoethyl ester (GSHE) or glutathione diethyl ester (GSHDE) afterward.^39^ We also synthesized glutathione (GSH)-capped gold nanoclusters using GSH instead of 3-MBA in the first synthetic procedure (Supporting Information Figure S2). The syntheses yielded uniform-sized gold nanoclusters with core diameters ranging from 1.3 to 2.7 nm (Figure 2a and Supporting Information Figure S2a). In this size range, the localized surface plasmon resonance (LSPR) band at the wavelength of 515 nm emerged above the core size of 2 nm, which is indicative of the transition from molecular to metallic nanoclusters (Figure 2b and Supporting Information Figure S2b).

**Figure 2.**
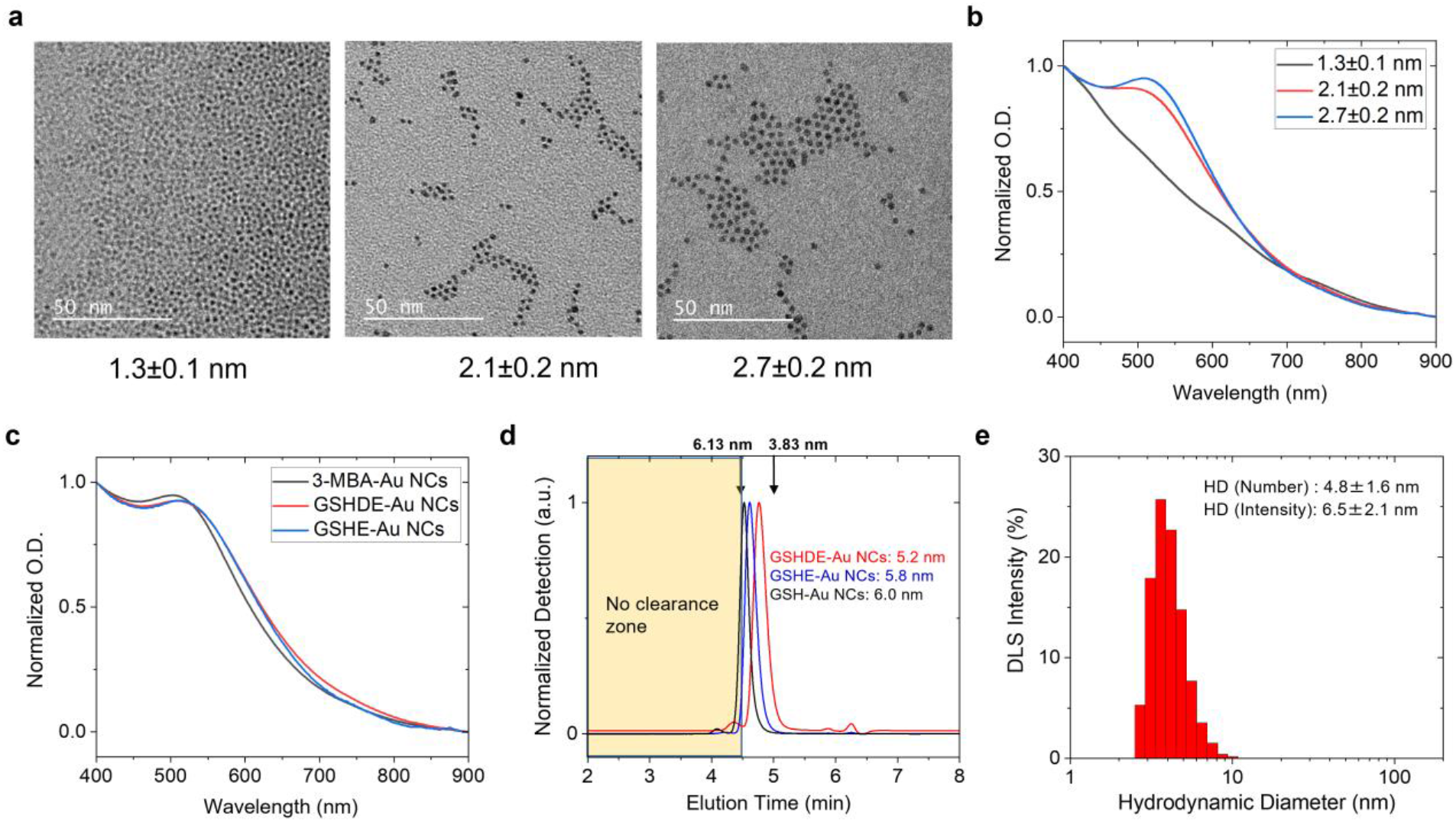
Synthesis of excretable gold nanocluster building blocks. (**a**) Transmission electron microscope (TEM) images of 3-mercaptobenzoic acid (3-MBA)-capped gold nanoclusters. Scale bar: 50 nm. (**b**) UV-VIS spectra of the gold nanoclusters displaying the emergence of localized surface plasmon resonance upon the size control over 2 nm. (**c**) UV-VIS spectra of the 2.7 nm gold nanoclusters before (3-MBA-Au NCs, black) and after ligand exchange with glutathione diethyl ester (GSHDE-Au NCs, red) and glutathione monoethyl ester (GSHE-Au NCs, blue). (**d**) Gel filtration chromatography of the glutathione (ester)-capped nanoclusters, showing that the hydrodynamic diameters (HDs) were controlled below the threshold of renal clearance. The size regime above the threshold is marked as yellow (see Supporting Information Figure S1 for standard calibration and Supporting Information Figure S2 for the information on glutathione-capped gold nanoclusters (GSH-Au NCs)). (**e**) Dynamic light scattering of the 2.7 nm GSHE-Au NCs exhibiting a similar HD as the size exclusion chromatography in (d).

Along with the precise control of core diameter, the compact glutathione (ester) ligands on the surface kept the metallic nanoclusters physiologically stable and their hydrodynamic diameter (HD) below the threshold for renal clearance (Figure 2c-e). The surface ligand exchange from 3-MBA to glutathione esters induced no significant change in the LSPR band, indicating that the metallic nanoclusters maintained their physiological stability without aggregation (Figure 2c). Previously, the supracluster approach utilized either hydrophobic or polymeric capping ligands in nanocluster building blocks that would induce poor dispersion in biological media or afford larger hydrodynamic size^31–34^. In contrast, gel filtration chromatography showed that the glutathione (ester)-capped 2.7 nm nanoclusters retained average HDs in the range from 5.2±0.6 nm to 6.0±0.7 nm in phosphate-buffered saline (Figure 2d, Supporting Information Figure S1, and Supporting Information Table S1). Dynamic light scattering (DLS) further confirmed that the average HD of the nanoclusters was comparable to the threshold for renal clearance of 6 nm (4.8±1.6 nm from number distribution and 6.5±2.1 nm from intensity distribution for the GSHE-capped gold nanoclusters, Figure 2e). We further confirmed their clearance capability by intravenously injecting the nanoclusters into nude mice and analyzing collected urine by inductively coupled plasma-mass spectrometry (ICP-MS), which revealed that 10∼20% of nanoclusters were rapidly eliminated through the renal clearance pathway within 2 days (Supporting Information Table S1). This result indicates that the synthesized nanoclusters are in the size range below the renal clearance cutoff.

Overall, we successfully produced metallic gold nanocluster building blocks with a size larger than the threshold to produce LSPR-generating metallic nanoclusters but smaller than the renal clearance cutoff. Note that the 2.7 nm core glutathione (ester)-capped gold nanoclusters have already reached the upper end of the renal clearance window below the HD of 6 nm. The results indicated that the size window for gold nanoclusters to meet all the criteria described above was very narrow within one nanometer (2∼3 nm).

### Synthesis and Raman spectroscopy of gold supraclusters

Next, we synthesized metallic supraclusters through the reaction of the glutathione (ester)-capped gold nanoclusters with bis(sulfosuccinimidyl) 2,2,4,4-glutarate (BS2G) crosslinker via the N-hydroxysuccinimide (NHS) coupling (Figure 3a). The supraclusters were synthesized with sizes and shapes varied upon the choice of the GSH, GSHE, and GSHDE-capped gold nanoclusters (GSH-Au, GSHE-Au, and GSHDE-Au nanoclusters, Figure 3b and c). In general, larger and more complex supraclusters were generated when the negative surface charge of the supraclusters was reduced by converting carboxylic acids in the ligand to esters from −46±3 mV (GSH-Au supraclusters) to −35±3 mV (GSHE-Au supraclusters) and −22±2 mV (GSHDE-Au supraclusters, Figure 3b and 3d). The GSH-Au and GSHE-Au nanoclusters produced supraclusters with spherical shapes with average diameters of 49(±9) nm and 192(±35) nm, respectively. When GSHDE-Au nanoclusters were used for synthesis, highly branched supraclusters with 249(±162) nm in length and 39(±10) nm in width were produced. The supraclusters formation resulted in a red shift of the LSPR wavelength from 515 nm (black, Figure 3e) to 555 nm (blue, Figure 3e) and a change in color of the colloidal solution from red to purple (inset, Figure 3e), which indicates plasmonic coupling of the nanocluster building blocks within the supraclusters.^31,40–43^ The LSPR wavelength was further red-shifted to 590 nm with broader band absorption at the NIR wavelength when more BS2G crosslinker was added in the supracluster synthesis (green, Figure 3e). Meanwhile, the DLS measurement revealed that it was accompanied by an increase in average size from 180 nm to 500 nm and broader size distribution with an increase in the polydispersity index (PDI) from 0.10 to 0.15 for the supraclusters (Figure 3f). When the synthetic conditions were adjusted to yield a PDI value of ∼0.1, the LSPR wavelength remained constant at 555 nm for all supraclusters of GSH-Au, GSHE-Au, and GSHDE-Au (Figure 3g), while their average HD ranged as 100 nm, 180 nm, and 370 nm, for the GSH-Au (black), GSHE-Au (blue), and GSHDE-Au supraclusters (red), respectively (Figure 3h).

**Figure 3.**
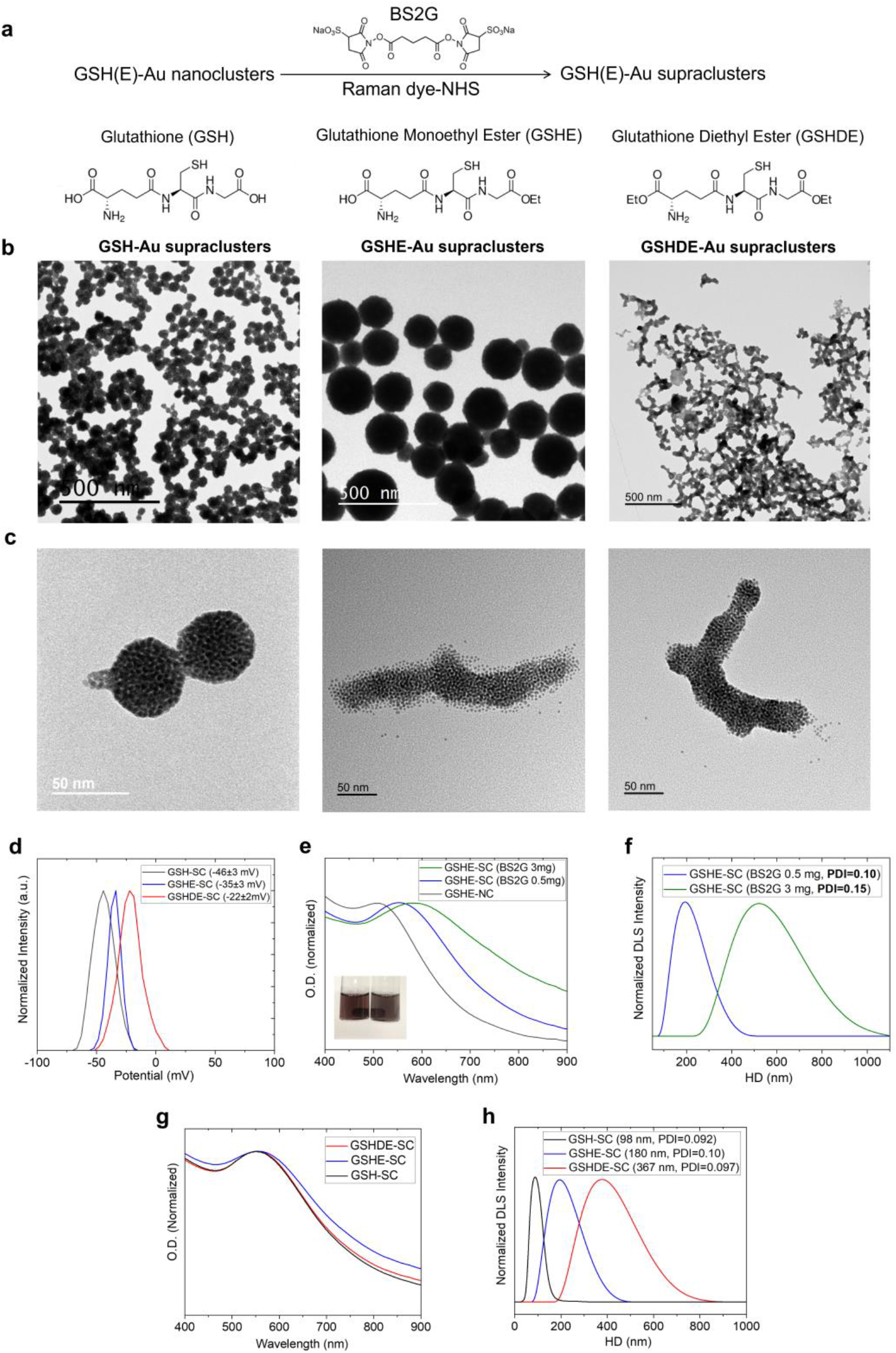
Synthesis of plasmonic supraclusters. (**a**) A reaction scheme for Raman supracluster synthesis. (**b**) Representative TEM images of supraclusters composed of glutathione-capped gold nanoclusters (GSH-Au supraclusters, left), glutathione monoethyl ester-capped gold nanoclusters (GSHE-Au supraclusters, middle), and glutathione diethyl ester-capped gold nanoclusters (GSHDE-Au supraclusters, right). (**c**) High-magnification images of spherical GSH-Au supraclusters (left) and branched GSHDE-Au supraclusters (middle and right). (**d**) Zeta potentials of GSH-Au supraclusters (black, GSH-Au SC), GSHE-Au supraclusters (blue, GSHE-Au SC), and GSHDE-Au supraclusters (GSHDE-Au SC, red). (**e**) UV-VIS-NIR spectra of the GSHE-Au nanoclusters (GSHE-Au NC, black) and GSHE-Au supraclusters (GSHE-Au SC, blue and green) exhibiting the localized surface plasmon resonance peak red-shifted with the increase of BS2G crosslinker from 0.5 mg (blue) to 3 mg (green) in the supracluster synthesis. The inset shows the color change of the colloidal solution before (left) and after supraclusters formation (right). (**f**) Dynamic light scattering (DLS) of the supraclusters displaying the average size and the PDI of the supraclusters increased with the increase of the BS2G crosslinker in the synthesis. (**g, h**) UV-VIS-NIR spectra (g) and dynamic light scattering (h) of the GSHDE-Au supraclusters (red, GSHDE-Au SC), GSHE-Au supraclusters (blue, GSHE-Au SC), and GSH-Au supraclusters (black, GSH-Au SC) when the size distribution was controlled with the PDI value of 0.1.

To make the supraclusters Raman scattering, we added NHS-functionalized NIR resonant Raman dyes during the supracluster synthesis to simultaneously proceed with their conjugation with the supraclusters. We tested the NIR non-fluorescent quencher molecules of SQ740 (SETA BioMedicals) and Tide Quencher™ 7WS (TQ7WS, AAT Bioquest) as Raman dyes to benefit from their NIR-absorption for resonant Raman scattering while reducing the fluorescence background (Figure 4 and Supporting Information Figure S3).

**Figure 4.**
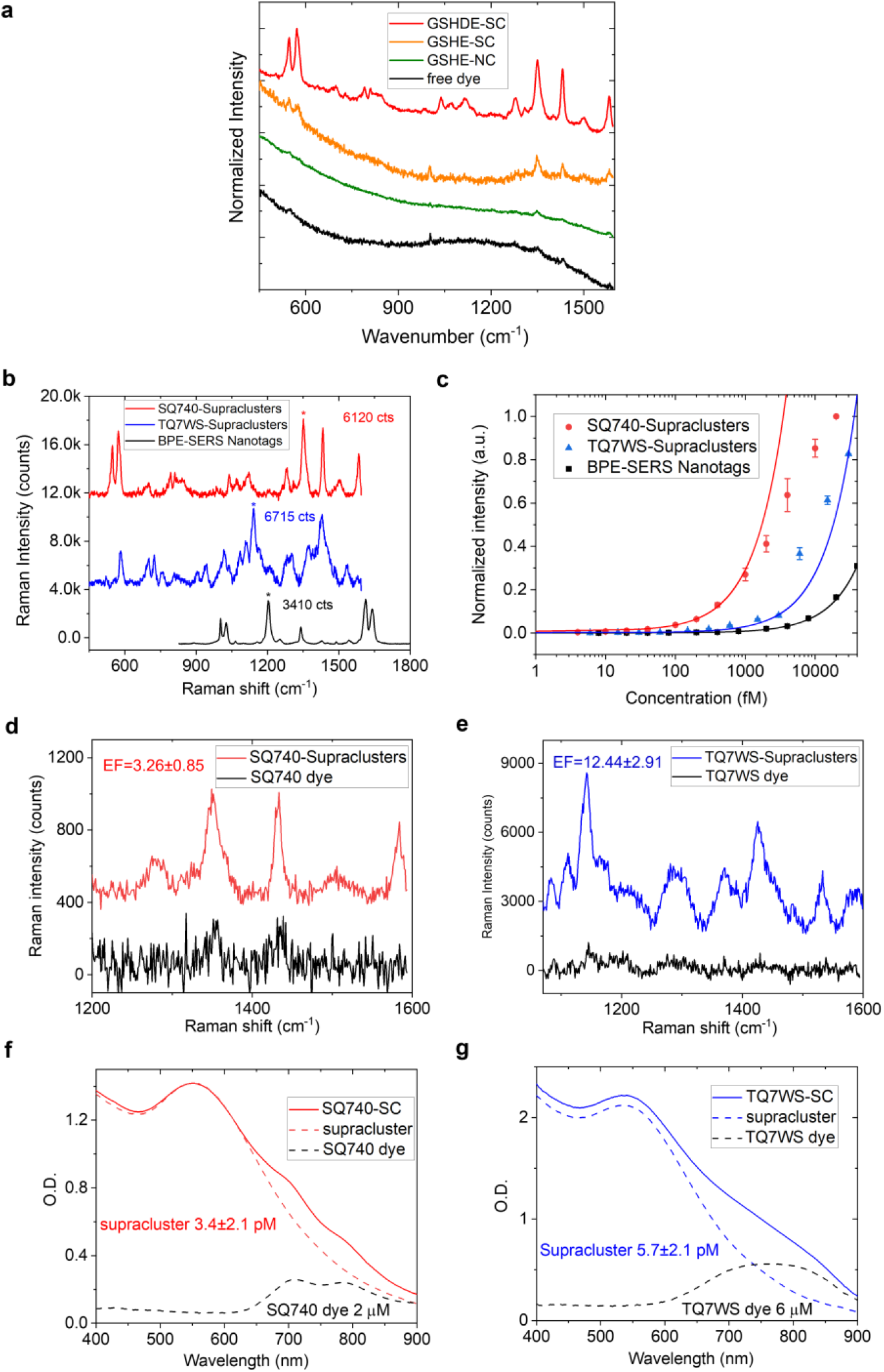
Raman spectroscopy of supraclusters. (**a**) The emission spectra of SQ740 Raman dyes in aqueous solution (black, free dye) and the SQ740-conjugated GSHE-Au nanoclusters (green, GSHE-NC), GSHE-Au supraclusters (orange, GSHE-Au SC), and GSHDE-Au supraclusters (red, GSHDE-Au SC). (**b**) Raman spectra of the SQ740-conjugated GSHDE-Au supraclusters (red, SQ740-Supraclusters), TQ7WS dyes-conjugated GSHDE-Au supraclusters (blue, TQ7WS-Supraclusters), and the SERS nanotags of 1,2-bis(4-pyridyl) ethylene (black, BPE-SERS Nanotags) at the same nanoparticle concentration 1 pM. (**c**) Normalized standard curves of integrated Raman intensities of the SQ740-Supraclusters (red), TQ7WS-Supraclusters (blue), and the BPE-SERS Nanotags (black). The limit of detections (LODs) calculated from the curves were 15.3±4.4 fM (red), 14.9±4.6 fM (blue), and 154±93 fM (black) concentration of the nanoparticles, respectively (see Supporting Information Figure S4). The Raman intensities in the plot was normalized to the integrated Raman intensity of SQ740-Supraclusters measured at 10 pM. (**d, e**) The estimation of the enhancement factor (EF) of the supraclusters. Comparison of Raman spectra of the SQ740-Supraclusters and free SQ740 Raman dyes in aqueous solution when the supracluster-loaded dye concentration was matched the same as the free Raman dyes (d). Comparison of Raman spectra of the TQ7WS-Supraclusters and free TQ7WS Raman dyes in aqueous solution when the supracluster-loaded dye concentration was matched the same as the free Raman dyes (e). (**f, g**) UV-VIS-NIR spectra of the SQ740-Supraclusters (SQ740-SC, f) and the TQ7WS-supraclusters (TQ7WS-SC, g), which were fitted with non-conjugated GSHDE-Au supraclusters to derive the loaded amount of the Raman dyes per supracluster. For Figures 4b to e, the fluorescence backgrounds were removed, and only the Raman components were taken into account. The error bars (c) represent the standard deviations of the Raman intensities collected from multiple points (n=300) scanning per measurement.

Despite the intrinsic fluorescence quenching effect of the Raman dyes, the free Raman dyes of SQ740 in an aqueous solution (black, Figure 4a) and the SQ740-conjugated GSHE-Au nanoclusters (green, Figure 4a) showed fluorescence background that was still dominant over Raman scattering when excited at 785 nm. However, when the Raman dyes were conjugated to the GSHE-Au supraclusters, the contribution from Raman scattering increased, and fingerprint-like Raman peaks became more prominent (orange, Figure 4a). It turned out that, among the supraclusters, the GSHDE-Au supraclusters displayed the most prominent Raman spectra of the Raman dyes with significant fluorescence suppression (red, Figure 4a).

We compared the Raman scattering brightness of the SQ740 (red, SQ740-Supraclusters, Figure 4b) and TQ7WS (blue, TQ7WS-Supraclusters, Figure 4b)-conjugated GSHDE-Au supraclusters to that of the commercially available 60 nm gold nanoparticles-based SERS nanotags of trans-1,2-bi-(4-pyridyl) ethylene (black, BPE-SERS Nanotags, Oxonica Materials, Inc, Figure 4b). Both supraclusters exhibited the Raman peak intensity that is twice as high as that of the BPE-SERS nanotags at 1 pM, and the brightness of the supraclusters, which was defined by the spectrally integrated Raman intensity, was one order of magnitude higher than the BPE-SERS nanotags (Figure 4b). The sensitivity and the limit of the detection for the Raman scattering brightness for the supraclusters were one order of magnitude higher (14.9±4.6 fM for TQ7WS-Supraclusters, blue, and 15.3±4.4 fM for SQ740-Supraclusters, red, Figure 4c and Supporting Information Figure S4) than the BPE-SERS nanotags (154±93 fM, black, Figure 4c and Supporting Information Figure S4), which demonstrates the potential for the supraclusters to replace large gold nanoparticles-based SERS nanotags for *in vivo* Raman imaging application.

To elucidate the origin of bright Raman scattering of the supraclusters, we first estimated the enhancement factor (EF) by comparing the Raman intensity when the concentration of the supraclusters-loaded dyes was matched with that of free dyes (Figure 4d and e). The enhancement factor determined for the supraclusters was 3.26±0.85 for the SQ740 dyes (Figure 4d) and 12.44±2.91 for the TQ7WS dyes (Figure 4e), which were not as large as that for the large gold nanoparticles-based SERS nanotags (∼10,000).^44^ Furthermore, no Raman signals were detected when the supraclusters were excited at the wavelength of 532 nm, which is in the resonance of the localized surface plasmon for the supraclusters, indicating that there was no significant contribution of the local electromagnetic field enhancement to the brightness of the supraclusters. Note that the size of the nanocluster building blocks of 2.7 nm, which was limited by the threshold for renal clearance, also limited the strength of plasmonic coupling.^45^ On the other hand, previous reports suggested that even slightly larger nanocluster building blocks over 5 nm allowed strong plasmonic coupling within the supraclusters that resulted in the broad SPR band at NIR, which would further increase the enhancement factor. However, this will sacrifice the clearance efficacy of the nanocluster building blocks.^37^

On the other hand, the GSHDE-Au supraclusters demonstrated ultrahigh loading capability of bright resonant Raman dyes and suppression of fluorescence background by the large metallic supraclusters. The number of Raman dyes loaded in the supraclusters was determined from the extinction spectra by fitting with the unconjugated GSHDE-Au supraclusters (Figure 4f and g), which yielded 5.9(±2.2)×10^5^ dyes (SQ740, Figure 4f) and 10.0(±2.8)×10^5^ dyes (TQ7WS, Figure 4g) per supracluster. This loading capacity outperformed the estimated maximum loading capacity of the BPE dyes on 90 nm gold nanoparticles (14,000 dyes per nanoparticle)^46^ by up to 70-fold. Furthermore, the large supraclusters provided a significant reduction of fluorescence background (×6) for the SQ740 supraclusters, which further enabled facile identification of the Raman spectral fingerprint (Figure 4a). We interpret the fluorescence quenching due to efficient charge transfer between the supraclusters and the Raman dyes.^47^ The fluorescence quenching benefits from the supracluster formation since larger nanoparticles allow a reservoir for efficient charge transfer with resonant dyes.^48^ Note that fluorescence quenching was most prominent for the GSHDE-Au supraclusters, which provided the most reduced negative surface charge among the supraclusters (Figure 3d) and would facilitate more efficient charge transfer.

Considering the Raman cross-section of NIR-resonant Raman dyes is typically ∼10^−25^ cm^2^/molecule, which is four orders of magnitude brighter than the non-resonant Raman dyes of BPE (∼10^−29^ cm^2^/molecule),^49,50^ overall, we elucidate that the contribution of high loading (×70) of bright NIR-resonant Raman dyes (×10^4^), a small enhancement factor (×10) along with fluorescence quenching by the large metallic supraclusters were sufficient to make the supraclusters as bright as the large gold nanoparticles-based SERS nanotags with large local electromagnetic field enhancement (×10^4^∼10^6^). This demonstration of SERS-compatible Raman scattering brightness of supraclusters is of significant importance since it enables combining the biodegradable design of supraclusters and the *in vivo* imaging utility of SERS imaging.

### *In vivo* Raman imaging of tumors

We also investigated the pH stability of the supraclusters within the range of pH 3.0 to 7.4. The supraclusters were stable in acidic conditions of pH ranging from 3 to 5.5 but disassembled into nanocluster building blocks at pH above 6, as shown in the transmission electron microscope (TEM) images and blue shift of the LSPR peak from 555 nm to 515 nm (Supporting Information Figure S5a and b). Along with the disassembly above pH 6.0, the Raman intensity was reduced by half, and the contribution of the fluorescence background to overall emission increased at pH 7.4 (Supporting Information Figure S5c). Such pH-responsive disassembly indicates imperfect covalent crosslinking between the nanoclusters, and we attribute the supracluster formation to both covalent crosslinking of the nanoclusters with both ends of BS2G and van der Waals interaction among the BS2G-conjugated nanoclusters on one side (Supporting Information Figure S6a). Then, the deprotonation of the carboxylic acid on the BS2G-conjugated nanoclusters at a higher pH would result in negative surface charge and repulsive forces between the nanoclusters for disassembly (Supporting Information Figure S6b).^51^

Since cancer provides acidic cellular pH and *in vivo* tumor microenvironment,^52^ we utilized the pH-response of the Raman supraclusters for *in vivo* imaging of the acidic tumor environment. We prepared 4T1 subcutaneous tumor xenograft models in nude mice, and 20 μL of 30 pM SQ740-supraclusters in saline solution were administered intratumorally (n=3) in the same way as the BPE-SERS nanotags that were previously utilized for *in vivo* Raman imaging after intratumoral injection.^15^ The subcutaneous tumors were scanned with a Raman microscope under 785 nm excitation while the mice were under anesthesia (Figure 5a). For a control experiment, we also performed *in vivo* Raman imaging of the normal tissue region where the supraclusters were administered subcutaneously.

**Figure 5.**
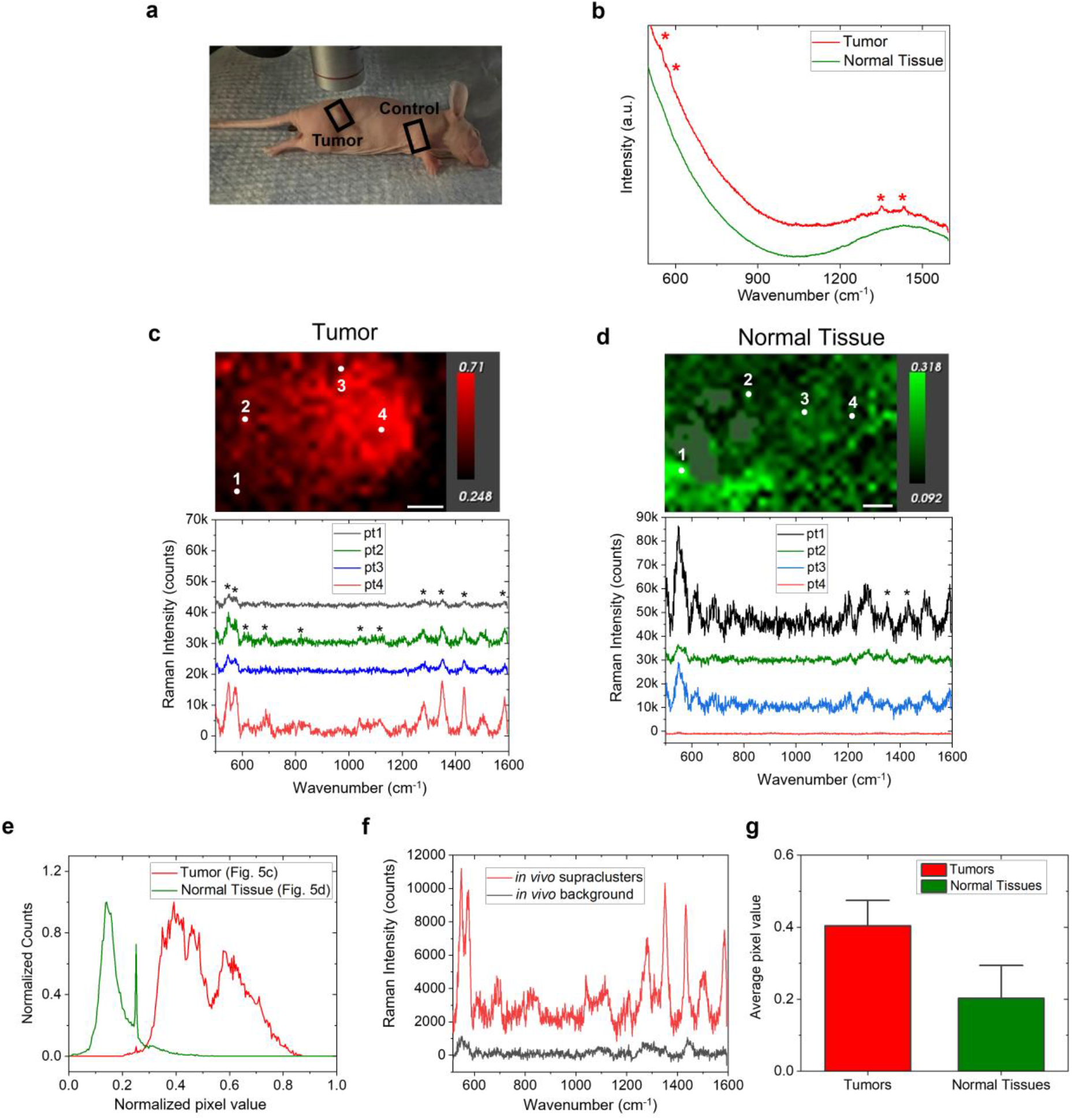
*In vivo* Raman imaging of supraclusters in tumors. (**a**) A schematic photo of *in vivo* Raman imaging of a tumor xenograft nude mouse under a Raman microscope. The regions of interest (ROIs) were defined as rectangles for the tumor site (tumor) and normal tissue (control). (**b**) The representative *in vivo* spectra after intratumoral (red) and subcutaneous (green) injections of the SQ740-Supraclusters into 4T1 tumor xenograft live nude mice. * indicate the spectral peaks corresponding to the SQ740-Supracluster Raman spectrum. (**c, d**) *In vivo* Raman images of supraclusters in the tumor site (c, tumor in a) and the normal tissue (d, control in a). The Raman images were generated through the least square analysis of the Raman spectra in each pixel to match with the pure Raman spectrum of the SQ740-supraclusters, followed by the normalization with mean center and scale to unit variance, the color-coding of the normalized intensities, and the visualization of the intensities corresponding to 5 to 95% of the distributions (upper panels). The pixel value closer to 1 means a higher correlation of the spectrum with the supracluster Raman spectrum. Scale bar: 2 mm. The representative four points Raman spectra were obtained from the Raman images in the upper panels (lower panels). * indicate the spectral peaks corresponding to the SQ740-Supracluster Raman spectrum. (**e**) The pixel intensity distributions in the Raman images of the tumor site (c) and the normal tissue region (d). (**f**) The representative Raman spectrum of tumors with (red) and without supraclusters injection (black). (**g**) The average pixel intensities extracted from the *in vivo* Raman images of tumors (red) and normal tissues (green) of the tumor xenograft nude mice (*P*=0.043, n=3). The error bars represent the standard deviations of the average pixel intensities collected from multiple mice.

Figure 5b shows the representative emission spectra of the tumors and normal tissue regions after the injections. While both regions displayed dominating fluorescence over Raman scattering, indicating that a significant amount of the supraclusters were dissociated, the Raman spectral peaks of the SQ740 supraclusters were clearly identified in tumors (red, Figure 5b).

The Raman images were generated through a least square analysis of the Raman spectra, applying the spectrum of the SQ740-supraclusters as a basis and displaying its correlation with the *in vivo* Raman spectrum at each pixel from 0 to 1. Figure 5c demonstrates that the Raman spectral image of a tumor exhibited a high correlation with the spectrum of the SQ740 supraclusters, with 5 to 95% of the pixel intensities to be within the range of 0.25∼0.71 with an average value of 0.50±0.12 (upper panel, Figure 5c and e). On the other hand, the Raman image of a normal tissue exhibited that the *in vivo* spectra match with the spectrum of supraclusters only within the intensity range of 0.09∼0.32 and with an average value of 0.17±0.06 (upper panel, Figure 5d and e). The representative *in vivo* Raman spectra of the tumor also shows that the Raman spectrum of the supraclusters dominantly contributed to the *in vivo* Raman spectra of the tumor, in which all the major peaks of SQ740 Raman spectrum can be identified (lower panel, Figure 5c). The average Raman intensity of the supraclusters at the tumor site was 11 times higher than that of the background *in vivo* Raman spectra at the tumor site without supraclusters administration, demonstrating the supraclusters provide sufficient sensitivity for *in vivo* tumor imaging (Figure 5f). In contrast, at the normal tissue region, the Raman intensity of the supraclusters decreased below that of the background Raman spectra (lower panel, Figure 5d). For *in vivo* Raman images obtained from the 4T1 tumor xenograft mouse models (n=3), the contribution of the Raman spectrum of the supraclusters to the *in vivo* Raman images reflected to the average pixel intensity was 2.2 times greater in tumors (0.44±0.17) than in normal tissues (0.20±0.04) (*P*=0.043, Figure 5g). The contrast of the Raman response in tumors and normal tissues (×2.20) was similar to the previously reported *in vivo* photoacoustic imaging with pH-sensitive “smart” gold nanomaterials composed of 10 nm gold nanoparticles (×1.96).^53^ Our results demonstrate that while the narrow pH window for the bright Raman scattering inevitably reduced the Raman intensity of the supraclusters at the physiological condition *in vivo*, it was sufficient enough to perform *in vivo* Raman imaging of tumors that provide more acidic environments.^52^

### Pharmacokinetics of the supraclusters

Finally, we evaluated the long-term pharmacokinetics of the gold supraclusters after systemic administration to nude mice. We intravenously administered the supraclusters at the dose of 160 μg gold (8 mg per kg body weight), which was equivalent to 6 fmol of the supraclusters (200 μL × 30 pM). For comparison, we also administered 60 nm PEGylated gold nanoparticles in the same dosage of gold to the control group of mice. Then, the biodistribution of supraclusters and nanoparticles in the heart, lung, liver, spleen, and kidney were tracked with inductively coupled plasma-optical emission spectroscopy (ICP-OES) and TEM (Figure 6 and Supporting Information Figures S7 to S9). The supraclusters and the nanoparticles were mostly accumulated in the liver, spleen, and kidney, which is in accordance with the previous reports on the pharmacokinetics of large nanoparticles.^22,30,36,37^ They were not detected in the heart and lung over the period of analysis.

**Figure 6.**
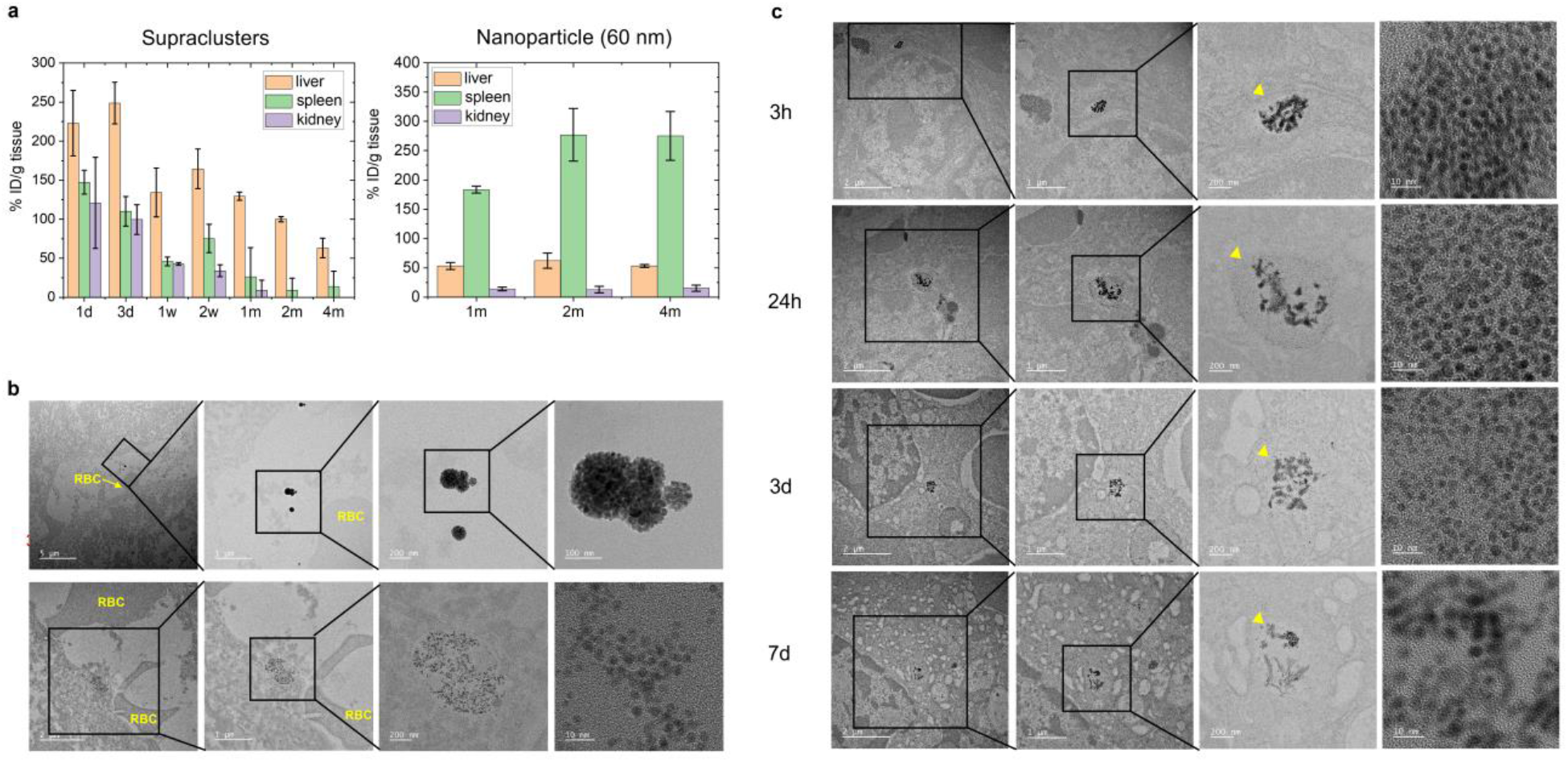
*In vivo* excretion of supraclusters. (**a**) *In vivo* biodistribution of the supraclusters (left) and nanoparticles (right) determined by quantifying gold ions in the liver, spleen, and kidneys with the ICP-OES (n=3). 160 μg of supraclusters or nanoparticles were intravenously administered, and the liver, spleen, and kidneys were collected at different time points. (**b**) TEM images of the GSHE-Au supraclusters (upper row) and dissociated nanoclusters (lower row) in the region of the liver near blood vessels. The regions identifying supraclusters or nanoclusters were highlighted as squares and enlarged in the next columns. (**c**) TEM images of supraclusters-uptaken spleens at 3 hours, 24 hours, 3 days, and 7 days after systematic administration of the GSHDE-Au supraclusters. The images were displayed in ascending order of magnification from left to right. The regions of supraclusters accumulation were highlighted as squares, enlarged in the next columns, and marked with yellow triangles in the third column. The fourth column is the high-resolution TEM images identifying dissociated nanoclusters.

While the 60 nm gold nanoparticles did not exhibit apparent signs of clearance from the liver and spleen, the supraclusters were excreted slowly over 4 months (Figure 6a). In the initial period of 1 and 3 days post-injection, TEM images revealed lots of supraclusters within the liver were located near blood vessels in either form of intact supraclusters (upper panel, Figure 6b) or dissociated nanoclusters (lower panel, Figure 6b). We also observed the supraclusters were internalized in the liver and spleen, dissociated into nanoclusters, and distributed through the tissues, resulting in a decrease in the local density of the dissociated gold nanoclusters in the tissues during the first 7 days after injection (Figure 6c and Supporting Information Figure S7). Such dissociation and redistribution of the nanoclusters were reflected in the rapid decrease of gold content within the tissues in the ICP-OES result (Figure 6a). After the first week, no significant change was observed in the TEM images (Supporting Information Figure S7), but the ICP-OES result showed slow and steady excretion of the gold content. As a result of the steady clearance, 75% (248 (3 days post-injection) to 63%ID/g) and 92% (147 (1 day post-injection) to 14%ID/g) of the initially internalized gold supraclusters were removed from the liver and spleen, respectively, over a period of 4 months. The level of gold excretion from the liver was superior to the previous results of metallic supraclusters (∼50%), which were designed to replace large gold nanoparticles.^36,37^ It is also notable that all the previously developed supraclusters demonstrated less efficient splenic than hepatic excretion^30^. In contrast, our supraclusters exhibit more efficient and nearly perfect excretion from the spleen. By far, little has been known about the splenic clearance mechanism of nanoparticles, which has been even less studied than hepatic excretion. We hypothesize that the physiologically stable glutathione esters-capped nanoclusters after the supracluster dissociation may undergo different splenic excretion mechanisms from either hydrophobic or physiologically unstable nanoclusters, which were previously used for supraclusters assembly. The biodistribution analysis also shows the complete clearance of the supraclusters from the kidneys within a month (Figure 6a). Compared to the liver and spleen, the TEM examination rarely identified the supraclusters in the kidneys but primarily at the glomerulus basement membrane and the intercellular space within proximal convoluted tubules as a form of dissociated nanoclusters, which suggests a contribution from the renal clearance pathway (Supporting Information Figure S8).

To achieve high excretion of the supraclusters, which is critical for the translation of this research, we precisely controlled the size of the nanocluster building blocks below the renal clearance threshold. It has not been determined to what extent the renal clearance of the nanoclusters contributes to the hepatic and splenic excretion of the supraclusters.^54^ However, it is clear that reducing the size of the nanocluster building blocks is also critical for faster hepatic clearance.^55^ Previous work on the excretion of 5 nm plasmonic nanoclusters-assembled supraclusters showed inefficient excretion compared to 1 nm molecular nanoclusters-assembled supraclusters.^37^ While a reduction in the size of the nanoclusters down to 1 nm led to more efficient excretion, this led to a loss of metallic properties for the gold nanoclusters. Overall, our result is the first to demonstrate significant excretion of metallic gold structures from all three RES organs of the liver, spleen, and kidney.

In contrast to the efficient dissociation and excretion of the supraclusters, the TEM images of the intravenously administered PEGylated 60 nm gold nanoparticles taken up by the liver and spleen displayed no morphological change over the same period of 4 months, which is in accordance with the previous reports (Supporting Information Figure S9).^22^

We examined the histopathology of the supraclusters-administered mice from the first day and week to the fourth month after the injection. The hematoxylin and eosin (H&E) stained tissues of the liver, spleen, kidney, heart, and lung revealed no evidence of inflammation and tissue damage at both the early and late stages of supracluster uptake (Supporting Information Figure S10). This result suggests good long-term *in vivo* biocompatibility of the supraclusters and further strengthens their translatability in addition to their efficient excretion.

## CONCLUSION

In conclusion, we have demonstrated gold supraclusters that show high excretion efficacy with greater than 75% hepatic excretion within 4 months and nearly complete clearance from the spleen and kidney while displaying prominent Raman scattering as bright as the large gold nanoparticle-based SERS nanotags. Therefore, this work lays a new strategy for designing gold nanomaterials for clinical applications. The demonstration of the *in vivo* Raman imaging described here further expands the translational utility of biodegradable gold supraparticles in addition to the previously reported computed tomography,^29,30^ photoacoustic imaging,^31,32^ and photothermal therapy.^33,34^ While the translation of *in vivo* Raman imaging has been attempted only in the colonoscopy setting since it can bypass the hepatic and renal clearance pathway,^56,57^ the Raman supracluster approach described here will broaden clinical applications beyond the gastrointestinal tract by enabling the systemic administration of gold nanomaterials.

## MATERIALS AND METHODS

### Synthesis of gold nanocluster building blocks

The gold nanoclusters were synthesized with the adoption and modification of the Kornberg method. To obtain glutathione monoethyl ester (GSHE, Cayman Chemical) and glutathione diethyl ester (GSHDE, BACHEM)-coated gold nanoclusters (GSHE-Au and GSHDE-Au nanoclusters), 3-mercaptobenzoic acid (3-MBA, Sigma-Aldrich)-capped gold nanoclusters (3-MBA-Au nanoclusters) were synthesized first. 0.047 g of gold chloride and 0.1295 g of 3-MBA were each dissolved in 10 mL of methanol. Then, the gold chloride solution was added to the 3-MBA solution, and 50 mL of deionized water and 6 mL of 1N sodium hydroxide solution were sequentially added, which produced a transparent yellow gold-mercaptobenzoic acid complex stock solution. The stock solution was held overnight, and its color changed to colorless transparent. 70 mL of methanol and 190 mL of deionized water were added to the overnight aged stock solution, and freshly dissolved 25 mg of sodium borohydride in 1 mL water was added to initiate the nanocluster growth. After 4.5 hours of reaction, 1.96 g of sodium chloride in 10 mL aqueous solution and 600 mL of methanol were added to precipitate the nanocluster powder overnight. The nanocluster powder was washed with methanol several times, dried in the air, and dispersed in 0.1M HEPES buffer (pH=7.3). For the ligand exchange of nanoclusters from 3-MBA to glutathione (ester), 100 mg of glutathione monoethyl ester or glutathione diethyl ester was added to the nanoclusters in HEPES buffer solution and reacted overnight. The ligand-exchanged nanoclusters were purified with a Microcon^(R)^ centrifugal filter (Millipore, molecular weight cutoff of 3kDa) and with PD-10 desalting columns. For control experiments, the size of the gold nanoclusters was controlled by changing the 3-MBA to gold chloride ratio from 3 to 7 and the added amount of 1N sodium hydroxide solution from 2 mL to 6 mL. For the synthesis of glutathione (GSH)-coated gold nanoclusters (GSH-Au nanoclusters), 0.082 g of gold chloride was dissolved in 10 mL of methanol, and 0.26 g of GSH was dissolved in 10 mL of deionized water, respectively, and the gold chloride in methanol solution was added to the GSH in water solution. Then, all the other synthetic procedures were kept the same as the 3-MBA-Au nanoclusters. All the nanoclusters were filtered with a 0.02 μm syringe filter (Whatman) before further use. For Supporting Information Figure S2, the average size of the GSH-Au nanoclusters was controlled by changing the molar ratio of gold chloride to GSH from 2 to 6. **Synthesis of Raman supraclusters**. For the synthesis of the Raman supraclusters, 1.4 mg (equivalent to 7 μmol gold ion and approximately 7 nmol of gold nanoclusters) of GSH, GSHE, and GSHDE-capped gold nanoclusters dispersed in 10 mL of 0.1M HEPES buffer (pH=7.3) were mixed with 1 mg of Bis[Sulfosuccinimidyl] glutarate (BS2G, Thermo Fisher Scientific) in 1 mL of 0.1M HEPES buffer solution and incubated at room temperature for 3 hrs. Then, 0.1 mg of NIR-resonant Raman dyes of SQ-740-NHS (SQ740, SETA Biomedicals) or Tide Quencher™ 7WS succinimidyl ester (T7WS, AAT Bioquest) was added to the incubating solution and held at room temperature overnight. After the overnight reaction, the supraclusters were retrieved via the centrifuge at 1,500g for 15 min, while unreacted nanoclusters remained as supernatant. The retrieved supraclusters were washed with deionized water four times.

### Characterizations of nanoclusters and supraclusters

The size and morphology of the gold nanoclusters and supraclusters were characterized using transmission electron microscopy (TEM, JEM 1400, JEOL, 120 kV with LaB6 emitter) and high-resolution TEM (FEI Tecnai G2 F20X-Twin microscope at 200 keV). With the TEM images, the size distribution of the nanoclusters was analyzed with ImageJ software (NIH). For hydrodynamic size measurements of the nanoclusters, Biozen SEC-2 size exclusion chromatography column (Phenomenex) was employed in the high-performance liquid chromatography (HPLC) setup, and the nanocluster size was derived by eluting 20.0 μL of the sample with PBS buffer at a flow rate of 0.5 mL/min and calibrating the chromatograph with that of the protein standards (Bio-Rad). The hydrodynamic size and zeta potential of supraclusters were determined using a Zetasizer NANO ZS-90 instrument (Malvern). The concentration of the supraclusters was determined using Nanoparticle Tracking Analysis (NanoSight, Malvern). The UV-VIS-NIR spectra of the nanoclusters and supraclusters were obtained using a 96-well microplate reader.

### Raman spectroscopy of supraclusters

The Raman spectra of the supraclusters were obtained by illuminating the Raman supraclusters solution in a 96-well plate using a confocal Raman microscope with 785 nm NIR-diode laser excitation (InVia Qontor, Renishaw), of which the incident power was 150 mW from the objective lens (×5, NA=0.15). The spectra were collected with an integration time of 1 second from the CCD spectrometer with a grating of 1200 groove/mm, which provides a spectral resolution of 1.07 cm^−1^. To obtain multiple numbers of spectra within the same solution for statistical analysis, the 96-well plate filled with the Raman supraclusters was scanned with the microscope with a step size from 100 to 200 μm to obtain 300 points per well.

The enhancement factor (*EF*) of the supraclusters was calculated using the following equation:

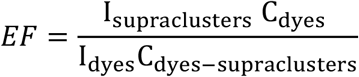

where I_supraclusters_ and C_dyes-supraclusters_ are the Raman intensity, and the concentration of the Raman dyes which were conjugated in supraclusters, respectively, and I_dyes_ and C_dyes_ are the Raman intensity and the concentration of the free Raman dyes in aqueous solution, respectively. C_dyes-supraclusters_ was determined by fitting the UV-VIS spectra of the Raman dyes-conjugated supraclusters with the unconjugated supraclusters to extract the Raman dye component. We hypothesized that the absorption coefficients of dyes within the supraclusters were the same as the free Raman dyes.

For the sensitivity measurement of the supraclusters, surface-enhanced Raman scattering (SERS) nanotags of S440 (Oxonica Materials, Inc.) were used as a control, in which *trans*-1,2-bis(4-pyridyl)-ethylene (BPE) was coated onto 60 nm gold nanoparticles and encapsulated with 35 nm-thick silica shell (BPE-SERS nanoparticles). The brightness of Raman scattering was determined from the area under the curve values of the Raman spectra by integrating the Raman intensity (*y*-axis) over wavenumber (*x*-axis). The limits of detection for the concentration of the supraclusters were calculated from the extrapolation of the sensitivity plots to the points, which intersect four times the standard deviation of the background Raman signals (Supporting Information Figure 4). **Animal experiments**. All procedures performed on the animals were approved by the Institutional Animal Care and Use Committee at Stanford University (APLAC #14465) and followed the NIH guidelines for the humane care of laboratory animals. Female 8-week-old nude mice (Envigo) were used for all *in vivo* studies.

### *In Vivo* Raman imaging of supraclusters in tumor xenograft nude mice

Subcutaneous tumor xenograft models were prepared by inoculating 4T1 breast cancer cells (2×10^6^ cells/mouse) into the hind leg of the nude mice (n= 3). We performed *in vivo* Raman imaging when the tumor size was grown to ∼1 cm^3^, as determined by caliper measurements.

For the *in vivo* Raman imaging, 20 μL of the 30 pM Raman supraclusters in saline solution was intratumorally administered into the subcutaneous tumors in the tumor xenograft nude mice (n=3). 1 hr after injection, the tumor-bearing mice were placed on the Raman microscope stage and put under anesthesia with 2∼3% isoflurane delivered in 100% oxygen for noninvasive Raman imaging (Figure 5a). The subcutaneous tumors were scanned with 785 nm diode laser equipped in the confocal Raman microscope, of which the incident power was 20∼80 mW from an objective lens (×5, NA=0.15), for 10 seconds duration time per 500 μm steps. The imaging generally took an hour to scan the entire area of a subcutaneous tumor.

The Raman images were generated through the least squares component analysis of the *in vivo* Raman spectrum at each pixel using the spectrum of the SQ740-supraclusters as a basis, normalization with a mean center, scale to unit variance, and color coding of the normalized intensities using WiRE 5.5 software (Figure 5c and d). The generated Raman images were processed with ImageJ for statistical analysis (Figure 5e and g).

### Biodistribution and animal study

160 μg of the supraclusters or nanoclusters were systematically administered to nude mice. As a control experiment, the same dose of 60 nm gold nanoparticles capped with mercapto-poly(ethylene glycol) was administered to the nude mice. Mice were sacrificed after 1, 3, 7, 14, 28 (1 month), 60 (2 months), and 120 (4 months) days, and liver, spleen, kidney, heart, and lung were collected and fixed with 4% paraformaldehyde. The organs were digested with 70% nitric acid, and the gold concentration was determined from inductively coupled plasma optical emission spectroscopy (ICP-OES). For the TEM analysis, the paraformaldehyde-fixed tissues were post-treated with 2% osmium for 40 min. The samples were dehydrated in a series of ethanol dilutions ranging from 50 to 100% and then kept in a series of absolute ethanol and Epon resin mixtures in ratios of 2:1, 1:1, and 1:2, followed by pure Epon resin overnight. The embedded resin was cured in a 60 °C oven for 48 hours. The samples were then cut into 80 nm ultrathin sections and mounted on a copper grid. The final samples were stained with uranyl acetate and lead citrate and examined under TEM (JEM-2100F, JEOL) at an accelerating voltage of 200 kV in the Korea Basic Science Institute, Chuncheon.

For the renal clearance test of the nanoclusters, 160 μg of the nanoclusters was injected into nude mice via tail vein. Then, the mice were placed in the metabolic cages. The urine was collected for 24 hrs and 48 hrs, dissolved in 70% nitric acid, and gold content was analyzed with inductively coupled plasma mass spectrometry (ICP-MS, n=4).

### Histological Analysis

Histological examination of the organs was performed based on the tissue sections of the liver, spleen, kidneys, heart, and lung, which were collected from the nude mice 1 day, 1 week and 4 months after the administration of 6 fmol of the supraclusters and stained with hematoxylin and eosin (H & E). A veterinary pathologist conducted the blind test for the H & E stained tissues.

### Statistical Analysis

The Raman intensity data were obtained from multiple points scanning measurements, and the mean and the standard deviation were derived from the multiple points data. The error bars in the normalized standard curves of the Raman intensities are the standard deviations (Figure 4c). The generated *in vivo* Raman images were processed with ImageJ software (NIH) for statistical analysis of the pixel intensities (Figures 5e and g). The standard deviation of average pixel intensities and their standard deviations were obtained from multiple measurements of three different tumor xenograft nude mice (n=3, Figure 5g). Unpaired *t*-test was performed to compare the two groups of tumors and normal tissues. Biodistribution of the supraclusters was derived from the ICP-OES analysis of the supraclusters-uptaken tissues with the errors bar representing the standard deviations of the gold content in organs measured from three different nude mice per period (n=3, Figure 6a).

## ASSOCIATED CONTENT

**Supporting Information**.

The Supporting Information is available free of charge online.

- Supporting Figures S1-S10 and Table S1. Detailed characterizations of the gold nanoclusters and supraclusters, pH-dependent properties, pharmacokinetics, and *in vivo* toxicity assessment of supraclusters.

## AUTHOR INFORMATION

### Author Contributions

J.H.Y., S.S.G., and J.R. conceived the project. J.H.Y. designed and carried out the experiments, analyzed the data, and wrote the manuscript. S.S.G. and J.R. analyzed the data, supervised the project, and co-wrote the manuscript. J.H.Y. and E.O.C. performed synthesis and characterizations of supraclusters. M.S.J. and S.-H.K. conducted TEM studies of the supraclusters-administered tissues. I.A., A.Z., and H.-Y.C. contributed to the animal study through nanoparticle administration and biodistribution analysis. S.T. and K.F. contributed to the animal study through tumor xenograft mice preparation. J.H.Y. performed *in vivo* Raman imaging of the tumor xenografts mice. J.H.Y., S.Y.L., and Y.I.P. conducted preparation and ICP-OES analysis of the supraclusters-administered tissues. The manuscript was revised through the contributions of all authors, and all the authors approved the final version of the manuscript.

### Notes

The authors declare no competing financial interests.

## ACKNOWLEDGMENT

We thank Dr. Kerriann Casey for the blind examination of the histology slides. The H&E slides of the mice organs were prepared through Animal Histology Services at the Department of Comparative Medicine at Stanford University. We thank Ji Yoon Kim in Korean Basic Science Institute, Gwangju Center, for her expertise with the ICP-OES analysis. This work was supported by the US National Institutes of Health (NIH) National Cancer Institute grant R01CA199656 (J.H.Y. and J.R.). This work is dedicated to the memory of Prof. Sanjiv Sam Gambhir (1962-2020), who led this project envisioning clinical translation of *in vivo* multiplexed Raman imaging and gold nanoparticles.

---

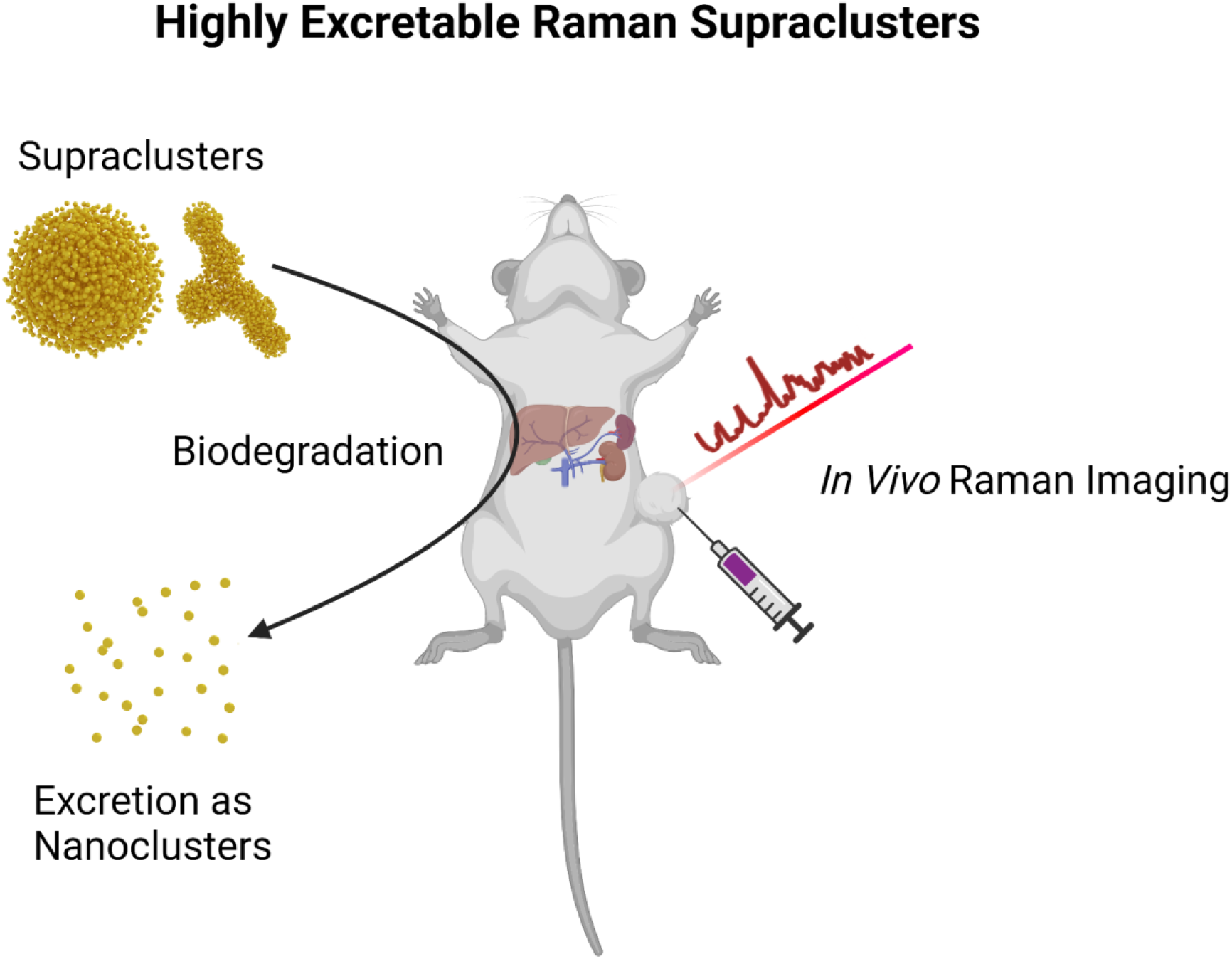

**Table of Contents**. A schematic of gold supraclusters that are biodegradable and excretable as nanoclusters and perform *in vivo* Raman imaging of tumors.

## Supporting Information

**Figure S1.**
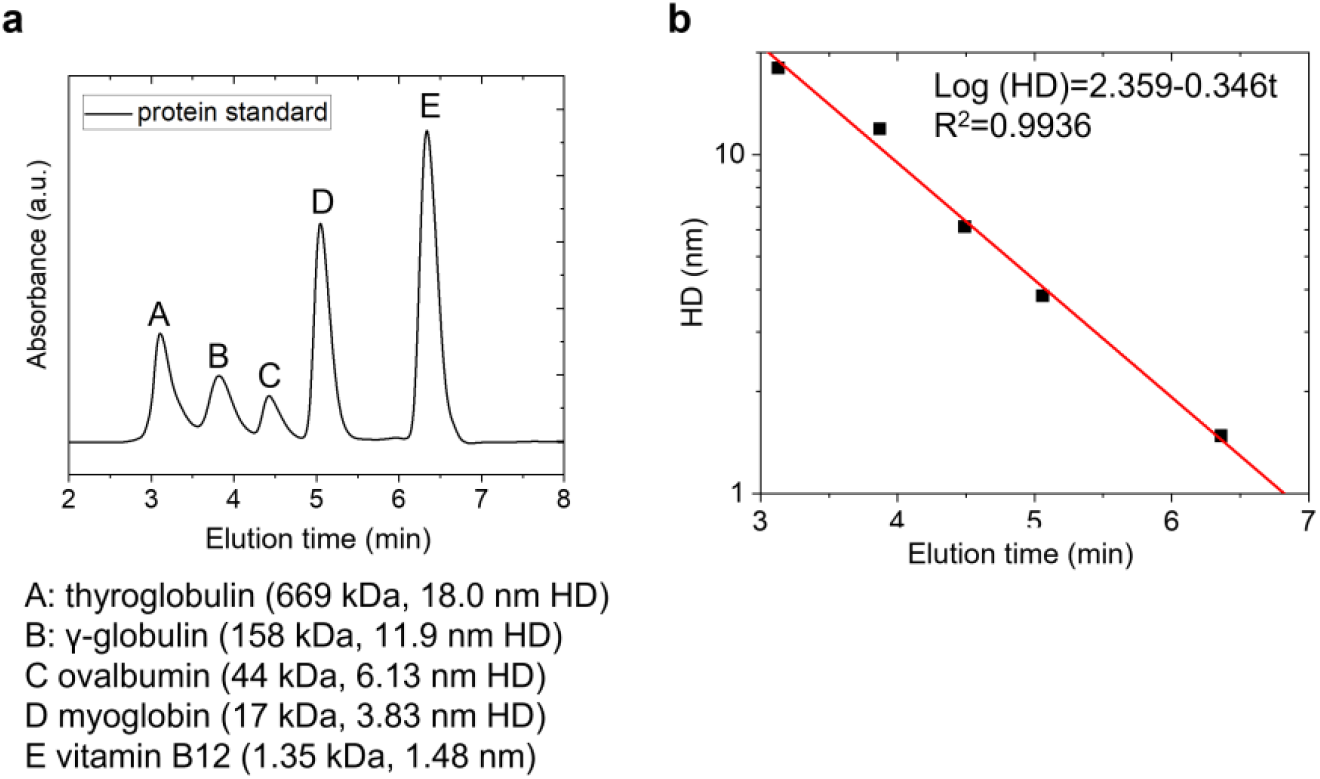
(**a**) Gel filtration chromatography of protein calibration standards. (**b**) Standard curve derived from (a) to estimate the hydrodynamic diameters (HDs) of the nanoclusters in Figure 2 (main text).

**Figure S2.**
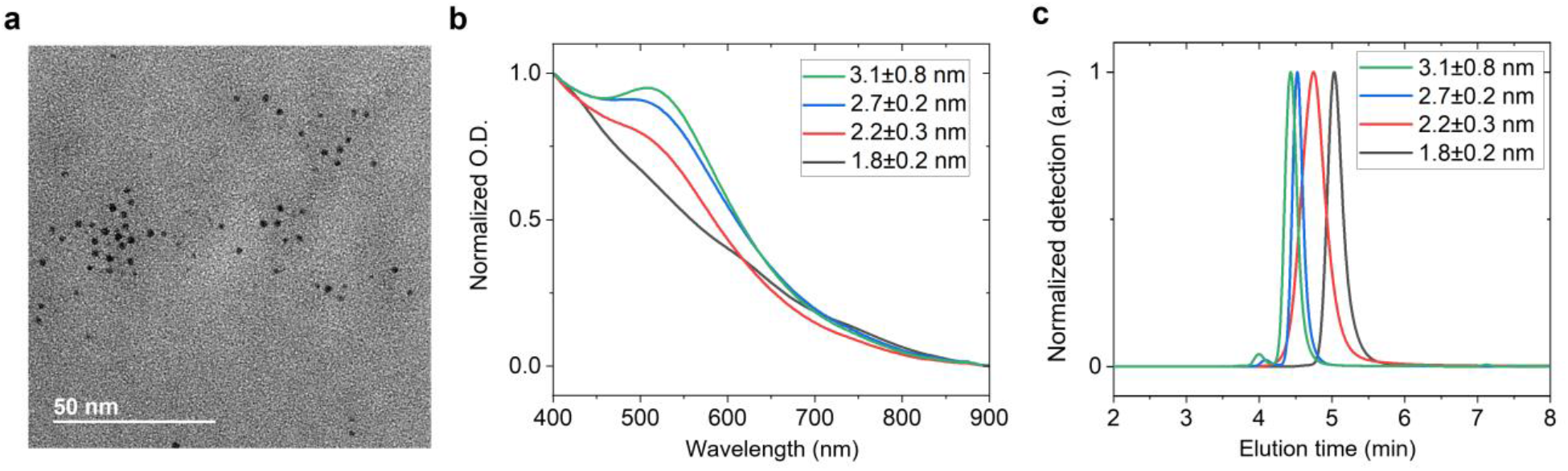
(**a**) A representative TEM image of 2.7±0.2 nm, glutathione-capped gold nanoclusters (GSH-Au NCs) utilized for supracluster synthesis in Figures 2 and 3 in the main text. (**b, c**) UV-VIS spectra (b) and gel filtration chromatography (c) of the GSH-Au NCs with their average size control from 1.8 nm to 3.1 nm. The localized surface plasmon resonance peak emerges when the average size of the nanoclusters increases over 2.2 nm (b). Gel filtration chromatography of the GSH-Au NCs shows that the hydrodynamic diameter (HD) varies with the core diameter changes below the threshold of renal clearance (c). Note that the size of the GSH-Au NCs was controlled by changing the relative ratio of glutathione to gold chloride from 2 (1.8 nm, black) to 6 (3.1 nm, green).

**Figure S3.**
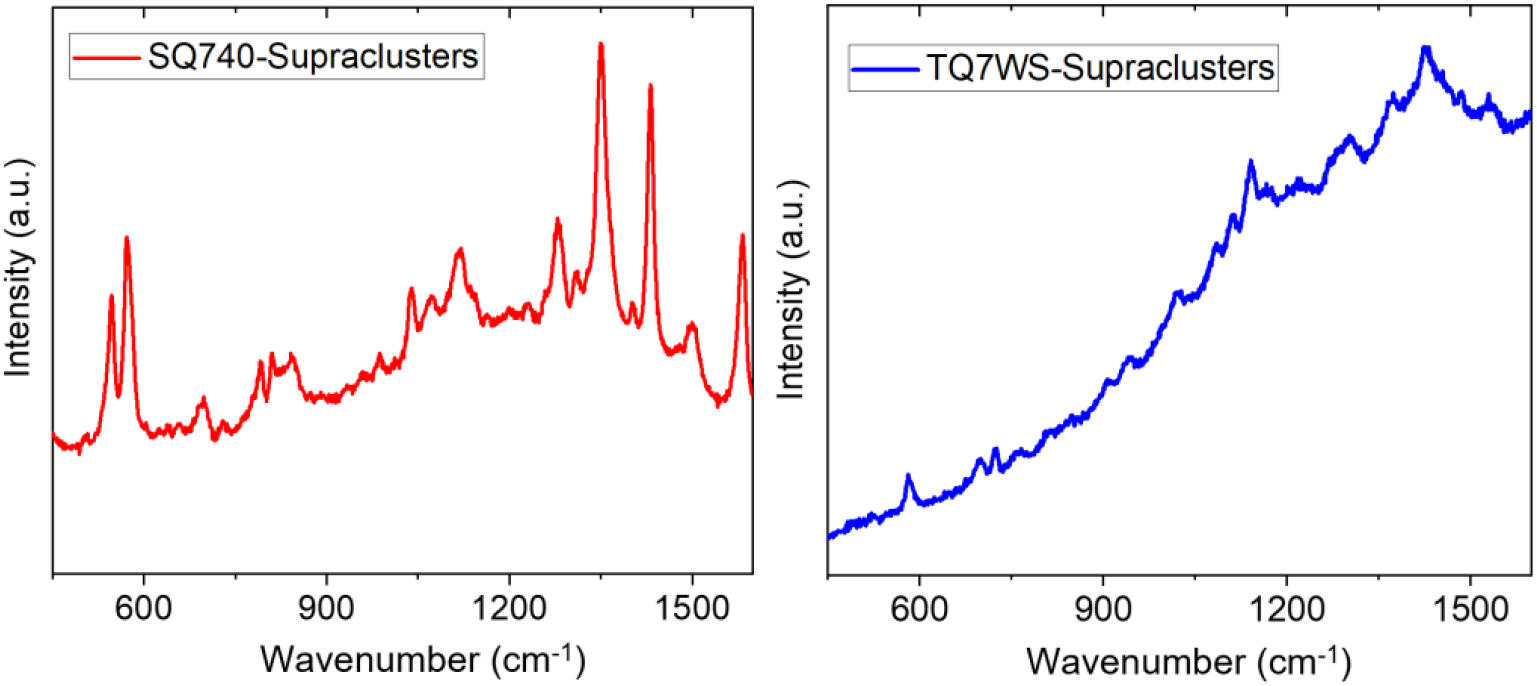
The emission spectra of SQ740-conjugated (red, left panel) and TQ7WS-conjugated (blue, right panel) GSHDE-Au supraclusters.

**Figure S4.**
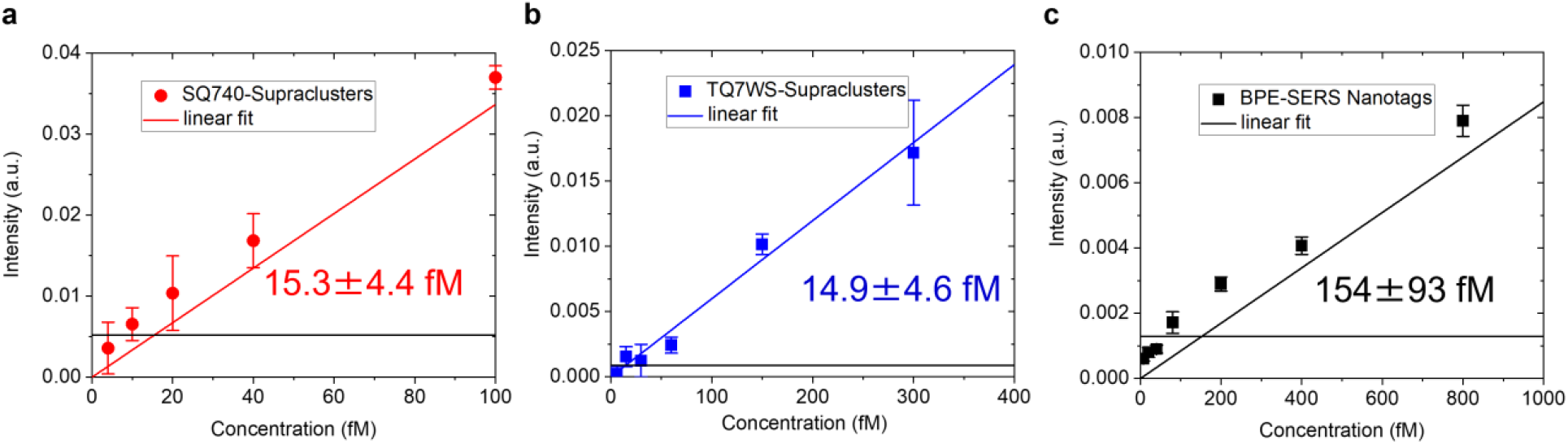
Limit of detections (LODs) calculated from the low concentration regime of the standard curves. (**a** to **c**) Linear plots of the Raman intensities at the low concentration regime (see Figure 4c in the main text) vs. concentration for the Raman scattering of the SQ740-Supraclusters (a), the TQ7WS-Supraclusters (b), and the BPE-SERS nanotags (c). The limits of detections (LODs) were calculated from the extrapolation of the linear plots to the points, which intersect four times the standard deviation of the background signals (horizontal line in each plot). The error bars represent standard deviations of the Raman intensities collected from multiple points (n=300) scanning per measurement.

**Figure S5.**
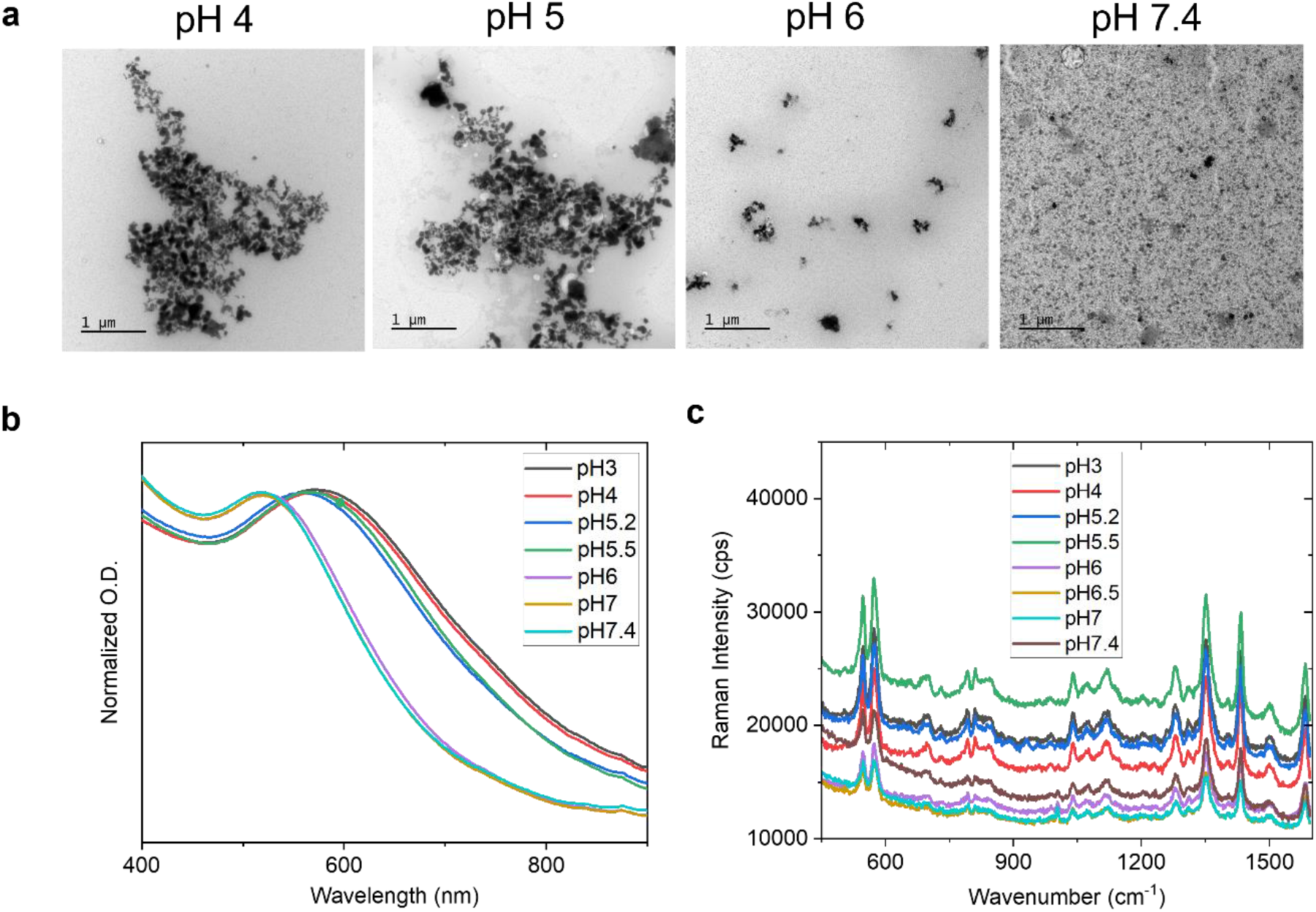
(**a**) TEM images of the GSHDE-Au supraclusters at pH 4, 5, 6, and 7.4. (**b**) UV-VIS-NIR spectra of the GSHDE-Au supraclusters under different pH from 3 to 7.4. The localized surface plasmon resonance (LSPR) peak shift from 555 nm to 515 nm at pH over 6 indicates dissociation of the supraclusters into the nanocluster building blocks. (**c**) Raman spectra of the SQ740-conjugated GSHDE supraclusters with the same pH variation as (b).

**Figure S6.**
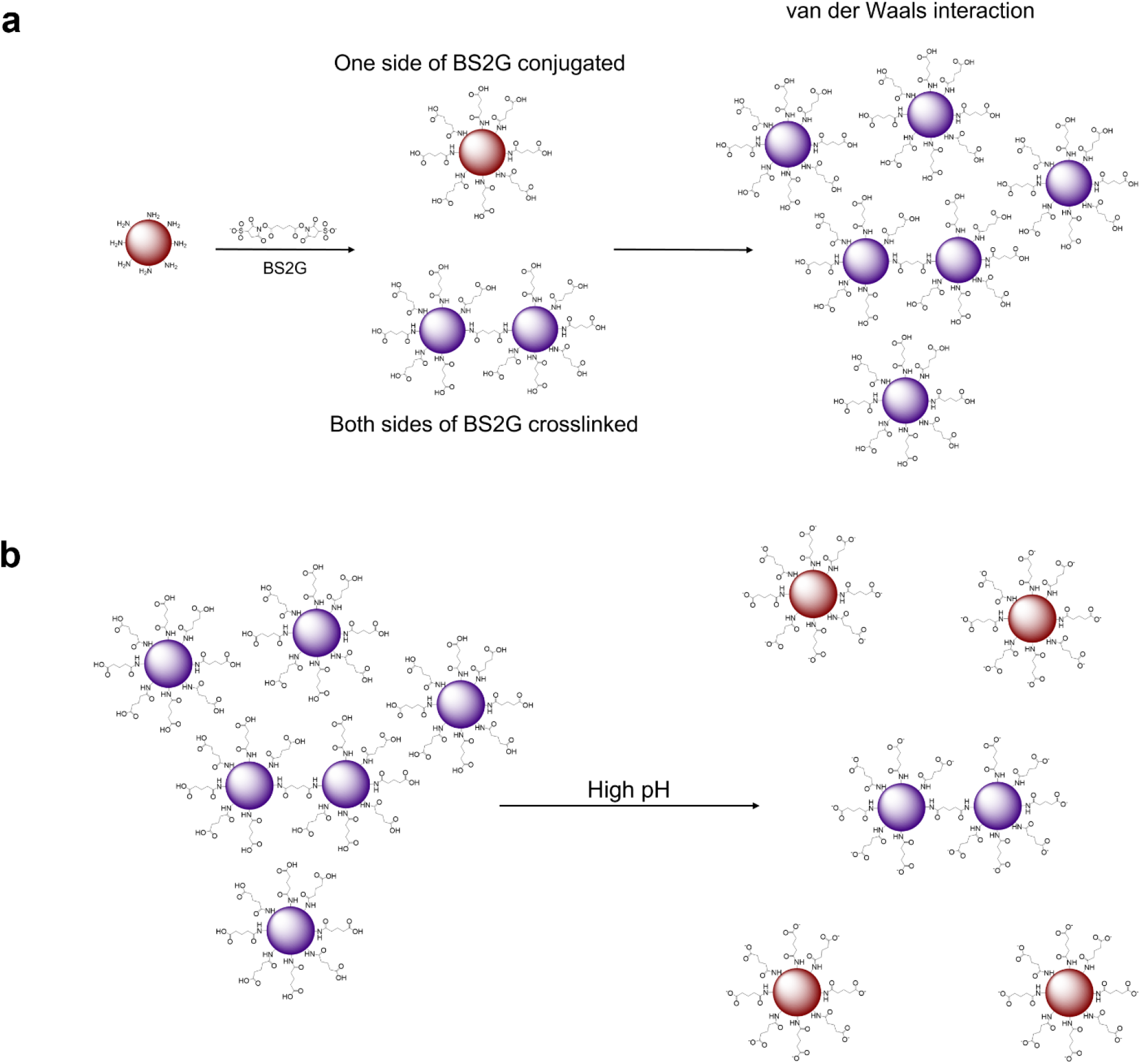
Schematics of the suggested mechanism of (**a**) supracluster formation and (**b**) disassembly at high pH.

**Figure S7.**
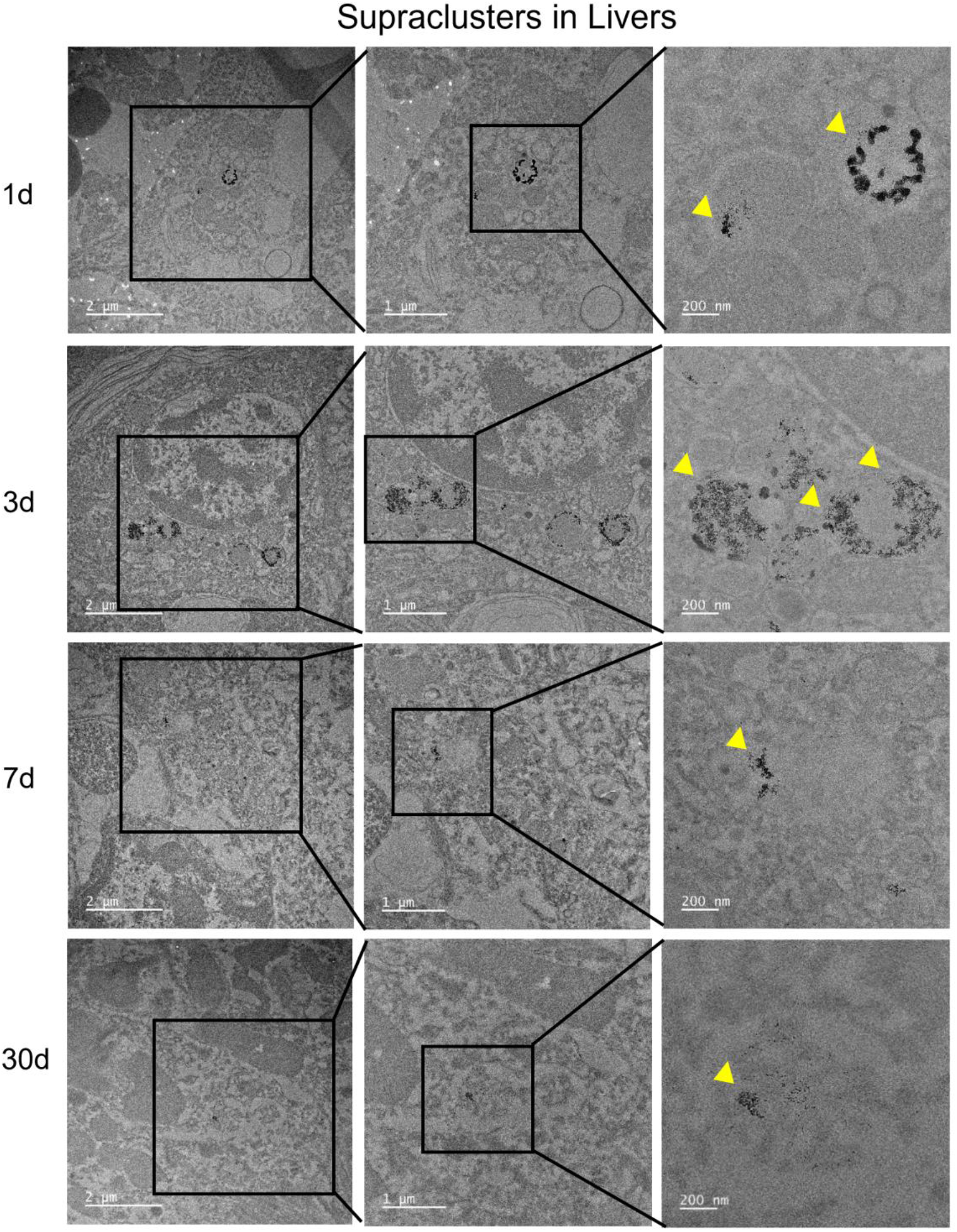
TEM images of supraclusters-uptaken livers at 1 day, 3 days, 7 days, and 30 days after systematic administration of the GSHDE supraclusters. The images were displayed in ascending order of magnification from left to right. The regions of supraclusters accumulation were highlighted as squares, enlarged in the next columns, and marked with yellow triangles in the third column.

**Figure S8.**
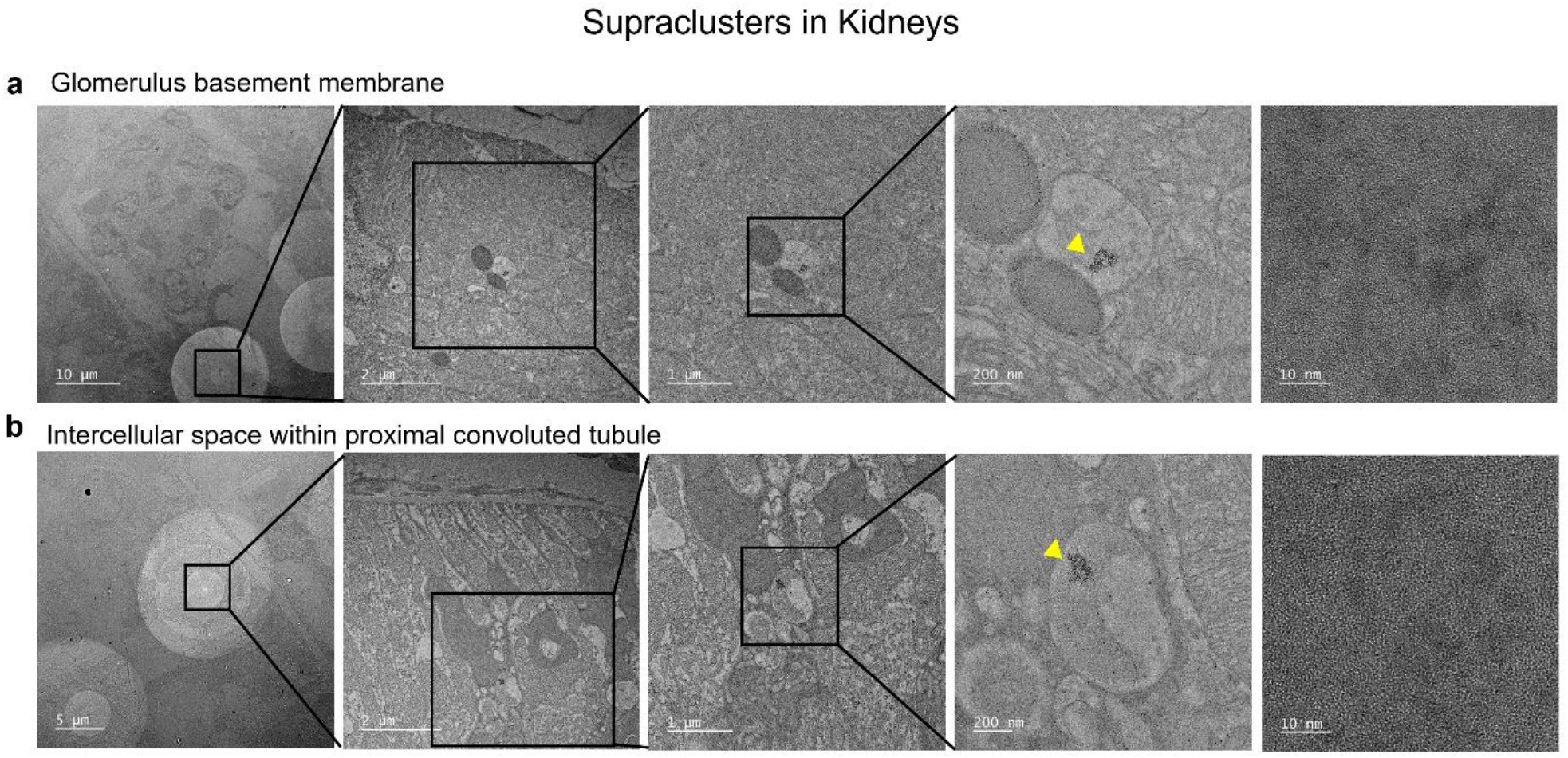
(**a, b**) TEM images of the dissociated nanoclusters which were found in the region of kidneys at the glomerulus basement membrane (a) and intercellular space within proximal convoluted tubule (b).

**Figure S9.**
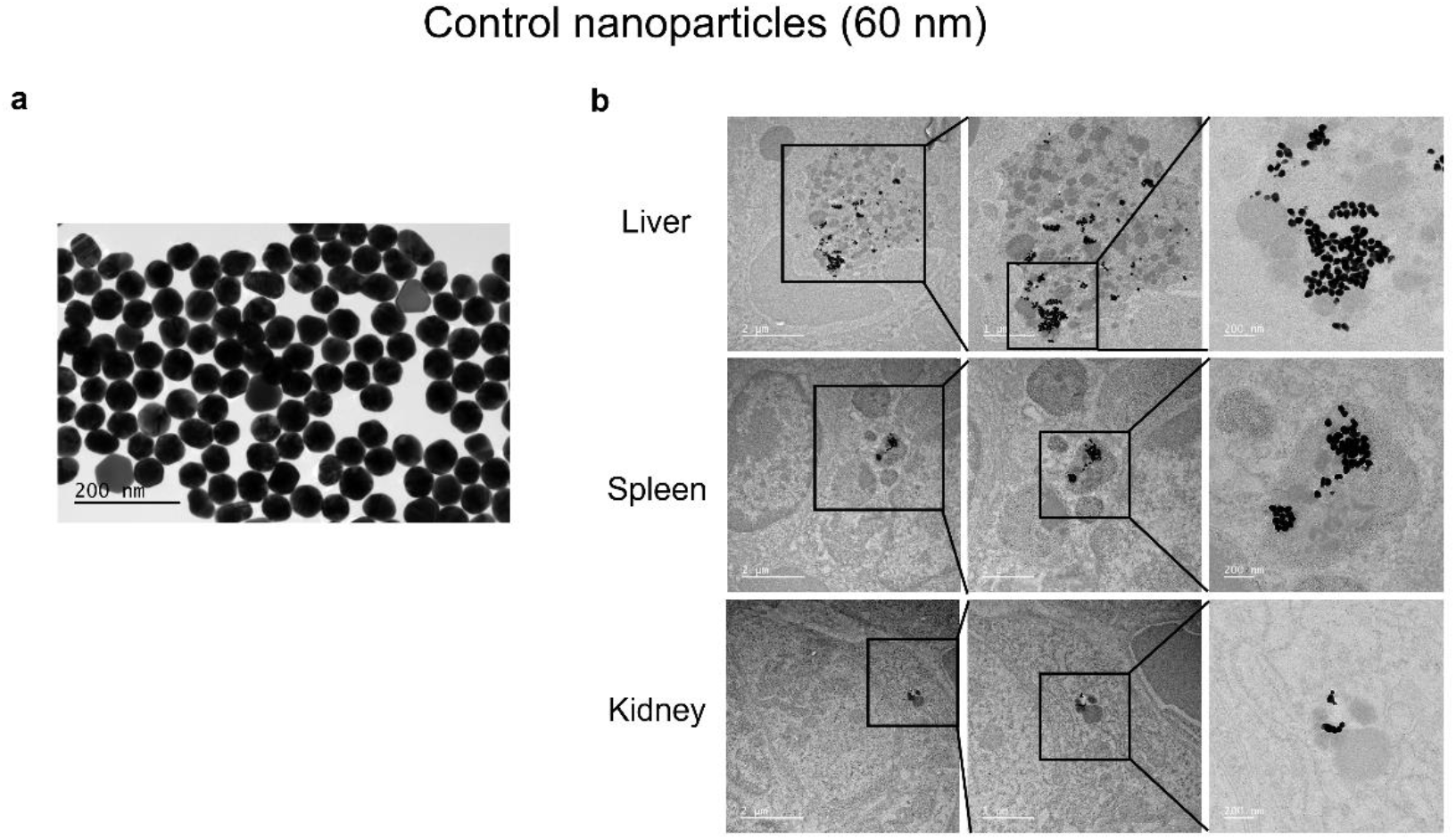
(**a**) TEM image of 60 nm PEGylated nanoparticles used as control. (**b**) TEM images of the liver, spleen, and kidney, 4 months after systematic administration of the gold nanoparticles. The regions identifying nanoparticles were highlighted as squares and enlarged in the next columns.

**Figure S10.**
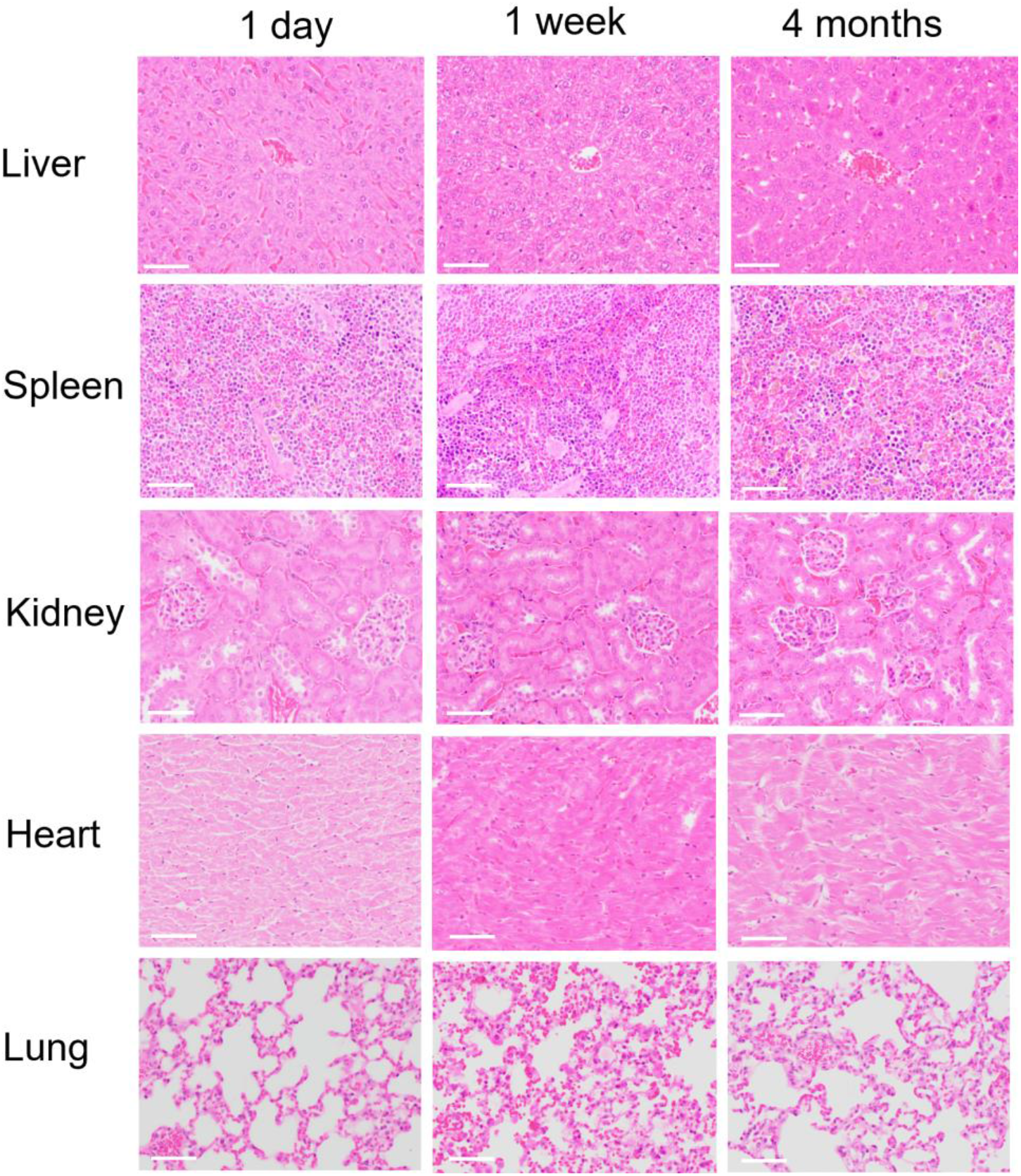
Histological examination of liver, spleen, kidney, heart, and lung tissues. treated with hematoxylin and eosin obtained at 1 day (left column), 1 week (middle column), and 4 months (right column) post-injection of supraclusters (*n* = 4). Scale bar: 50 μm. There is some red pigment within portions of the spleen, which is suspected as hemosiderin, a normal breakdown product of red blood cells. Overall, the examination confirms that there are no gross damages, inflammation, or necrosis of those organs particularly related to the administered supraclusters.

**Table S1.**
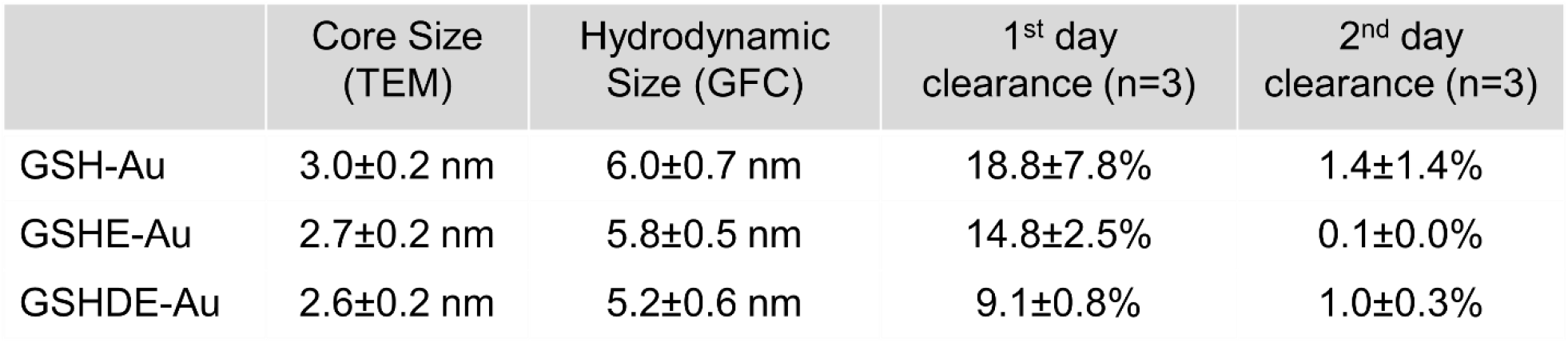
Core diameter, hydrodynamic diameter, and renal clearance efficiency on the first and second day after injection for the GSH-Au, GSHE-Au, and GSHDE-Au nanoclusters used for Raman supracluster synthesis.

|  | Core Size<br>(TEM) | Hydrodynamic<br>Size (GFC) | 1<sup>st</sup> day<br>clearance (n=3) | 2<sup>nd</sup> day<br>clearance (n=3) |
| --- | --- | --- | --- | --- |
| GSH-Au | 3.0±0.2 nm | 6.0±0.7 nm | 18.8±7.8% | 1.4±1.4% |
| GSHE-Au | 2.7±0.2 nm | 5.8±0.5 nm | 14.8±2.5% | 0.1±0.0% |
| GSHDE-Au | 2.6±0.2 nm | 5.2±0.6 nm | 9.1±0.8% | 1.0±0.3% |

## REFERENCES

(1) Keren, S.; Zavaleta, C.; Cheng, Z.; De La Zerda, A.; Gheysens, O.; Gambhir, S. S. Noninvasive Molecular Imaging of Small Living Subjects Using Raman Spectroscopy. Proc. Natl. Acad. Sci. U. S. A. 2008, 105, 5844–5849.

(2) Cheng, J.-X.; Xie, X. S. Vibrational Spectroscopic Imaging of Living Systems: An Emerging Platform for Biology and Medicine. Science. 2015, 350, aaa8870.

(3) Zhao, Z.; Shen, Y.; Hu, F.; Min, W. Applications of Vibrational Tags in Biological Imaging by Raman Microscopy. Analyst 2017, 142, 4018–4029.

(4) Mahadevan-Jansen, A.; Richards-Kortum, R. R. Raman Spectroscopy for the Detection of Cancers and Precancers. J. Biomed. Opt. 1996, 1, 31–70.

(5) Cordero, E.; Latka, I.; Matthäus, C.; Schie, I. W.; Popp, J. In-Vivo Raman Spectroscopy: From Basics to Applications. J. Biomed. Opt. 2018, 23, 71210.

(6) Jermyn, M.; Mok, K.; Mercier, J.; Desroches, J.; Pichette, J.; Saint-Arnaud, K.; Bernstein, L.; Guiot, M.-C.; Petrecca, K.; Leblond, F. Intraoperative Brain Cancer Detection with Raman Spectroscopy in Humans. Sci. Transl. Med. 2015, 7, 274ra19–274ra19.

(7) Huang, R.; Harmsen, S.; Samii, J. M.; Karabeber, H.; Pitter, K. L.; Holland, E. C.; Kircher, M. F. High Precision Imaging of Microscopic Spread of Glioblastoma with a Targeted Ultrasensitive SERRS Molecular Imaging Probe. Theranostics 2016, 6, 1075–1084.

(8) Nicolson, F.; Andreiuk, B.; Andreou, C.; Hsu, H.-T.; Rudder, S.; Kircher, M. F. Non-Invasive In Vivo Imaging of Cancer Using Surface-Enhanced Spatially Offset Raman Spectroscopy (SESORS). Theranostics 2019, 9, 5899–5913.

(9) Harmsen, S.; Huang, R.; Wall, M. A.; Karabeber, H.; Samii, J. M.; Spaliviero, M.; White, J. R.; Monette, S.; O’Connor, R.; Pitter, K. L.; Sastra, S. A.; Saborowski, M.; Holland, E. C.; Singer, S.; Olive, K. P.; Lowe, S. W.; Blasberg, R. G.; Kircher, M. F. Surface-Enhanced Resonance Raman Scattering Nanostars for High-Precision Cancer Imaging. Sci. Transl. Med. 2015, 7, 271ra7–271ra7.

(10) Andreou, C.; Neuschmelting, V.; Tschaharganeh, D. F.; Huang, C. H.; Oseledchyk, A.; Iacono, P.; Karabeber, H.; Colen, R. R.; Mannelli, L.; Lowe, S. W.; Kircher, M. F. Imaging of Liver Tumors Using Surface-Enhanced Raman Scattering Nanoparticles. ACS Nano 2016, 10, 5015–5026.

(11) Strobbia, P.; Cupil-Garcia, V.; Crawford, B. M.; Fales, A. M.; Pfefer, T. J.; Liu, Y.; Maiwald, M.; Sumpf, B.; Vo-Dinh, T. Accurate in Vivo Tumor Detection Using Plasmonic-Enhanced Shifted-Excitation Raman Difference Spectroscopy (SERDS). Theranostics 2021, 11, 4090–4102.

(12) Qiu, Y.; Zhang, Y.; Li, M.; Chen, G.; Fan, C.; Cui, K.; Wan, J.-B.; Han, A.; Ye, J.; Xiao, Z. Intraoperative Detection and Eradication of Residual Microtumors with Gap-Enhanced Raman Tags. ACS Nano 2018, 12, 7974–7985.

(13) Zavaleta, C. L.; Smith, B. R.; Walton, I.; Doering, W.; Davis, G.; Shojaei, B.; Natan, M. J.; Gambhir, S. S. Multiplexed Imaging of Surface Enhanced Raman Scattering Nanotags in Living Mice Using Noninvasive Raman Spectroscopy. Proc. Natl. Acad. Sci. 2009, 106, 13511–13516.

(14) Yu, J. H.; Steinberg, I.; Davis, R. M.; Malkovskiy, A. V; Zlitni, A.; Radzyminski, R. K.; Jung, K. O.; Chung, D. T.; Curet, L. D.; D’Souza, A. L.; Chang, E.; Rosenberg, J.; Campbell, J.; Frostig, H.; Park, S.; Pratx, G.; Levin, C.; Gambhir, S. S. Noninvasive and Highly Multiplexed Five-Color Tumor Imaging of Multicore Near-Infrared Resonant Surface-Enhanced Raman Nanoparticles In Vivo. ACS Nano 2021, 15, 19956–19969.

(15) Dinish, U. S.; Balasundaram, G.; Chang, Y.-T.; Olivo, M. Actively Targeted In Vivo Multiplex Detection of Intrinsic Cancer Biomarkers Using Biocompatible SERS Nanotags. Sci. Rep. 2014, 4, 4075.

(16) Ou, Y.-C.; Wen, X.; Johnson, C. A.; Shae, D.; Ayala, O. D.; Webb, J. A.; Lin, E. C.; DeLapp, R. C.; Boyd, K. L.; Richmond, A.; Mahadevan-Jansen, A.; Rafat, M.; Wilson, J. T.; Balko, J. M.; Tantawy, M. N.; Vilgelm, A. E.; Bardhan, R. Multimodal Multiplexed Immunoimaging with Nanostars to Detect Multiple Immunomarkers and Monitor Response to Immunotherapies. ACS Nano 2020, 14, 651–663.

(17) Langer, J.; Jimenez de Aberasturi, D.; Aizpurua, J.; Alvarez-Puebla, R. A.; Auguié, B.; Baumberg, J. J.; Bazan, G. C.; Bell, S. E. J.; Boisen, A.; Brolo, A. G.; Choo, J.; Cialla-May, D.; Deckert, V.; Fabris, L.; Faulds, K.; García de Abajo, F. J.; Goodacre, R.; Graham, D.; Haes, A. J.; Haynes, C. L.; Huck, C.; Itoh, T.; Käll, M.; Kneipp, J.; Kotov, N. A.; Kuang, H.; Le Ru, E. C.; Lee, H. K.; Li, J.-F.; Ling, X. Y.; Maier, S. A.; Mayerhöfer, T.; Moskovits, M.; Murakoshi, K.; Nam, J.-M.; Nie, S.; Ozaki, Y.; Pastoriza-Santos, I.; Perez-Juste, J.; Popp, J.; Pucci, A.; Reich, S.; Ren, B.; Schatz, G. C.; Shegai, T.; Schlücker, S.; Tay, L.-L.; Thomas, K. G.; Tian, Z.-Q.; Van Duyne, R. P.; Vo-Dinh, T.; Wang, Y.; Willets, K. A.; Xu, C.; Xu, H.; Xu, Y.; Yamamoto, Y. S.; Zhao, B.; Liz-Marzán, L. M. Present and Future of Surface-Enhanced Raman Scattering. ACS Nano 2020, 14, 28–117.

(18) Lane, L. A.; Qian, X.; Nie, S. SERS Nanoparticles in Medicine: From Label-Free Detection to Spectroscopic Tagging. Chem. Rev. 2015, 115, 10489–10529.

(19) Laing, S.; Jamieson, L. E.; Faulds, K.; Graham, D. Surface-Enhanced Raman Spectroscopy for in Vivo Biosensing. Nat. Rev. Chem. 2017, 1, 60.

(20) Hong, S.; Li, X. Optimal Size of Gold Nanoparticles for Surface-Enhanced Raman Spectroscopy under Different Conditions. J. Nanomater. 2013, 2013, 790323.

(21) Ali, M. R. K.; Rahman, M. A.; Wu, Y.; Han, T.; Peng, X.; Mackey, M. A.; Wang, D.; Shin, H. J.; Chen, Z. G.; Xiao, H.; Wu, R.; Tang, Y.; Shin, D. M.; El-Sayed, M. A. Efficacy, Long-Term Toxicity, and Mechanistic Studies of Gold Nanorods Photothermal Therapy of Cancer in Xenograft Mice. Proc. Natl. Acad. Sci. 2017, 114, E3110–E3118.

(22) Thakor, A. S.; Luong, R.; Paulmurugan, R.; Lin, F. I.; Kempen, P.; Zavaleta, C.; Chu, P.; Massoud, T. F.; Sinclair, R.; Gambhir, S. S. The Fate and Toxicity of Raman-Active Silica-Gold Nanoparticles in Mice. Sci. Transl. Med. 2011, 3, 79ra33.

(23) Falagan-Lotsch, P.; Grzincic, E. M.; Murphy, C. J. One Low-Dose Exposure of Gold Nanoparticles Induces Long-Term Changes in Human Cells. Proc. Natl. Acad. Sci. 2016, 113, 13318–13323.

(24) Zhou, C.; Long, M.; Qin, Y.; Sun, X.; Zheng, J. Luminescent Gold Nanoparticles with Efficient Renal Clearance. Angew. Chem. Int. Ed. Engl. 2011, 50, 3168–3172.

(25) Du, B.; Jiang, X.; Das, A.; Zhou, Q.; Yu, M.; Jin, R.; Zheng, J. Glomerular Barrier Behaves as an Atomically Precise Bandpass Filter in a Sub-Nanometre Regime. Nat. Nanotechnol. | 2017, 12, 1096–1102.

(26) Loynachan, C. N.; Soleimany, A. P.; Dudani, J. S.; Lin, Y.; Najer, A.; Bekdemir, A.; Chen, Q.; Bhatia, S. N.; Stevens, M. M. Renal Clearable Catalytic Gold Nanoclusters for in Vivo Disease Monitoring. Nat. Nanotechnol. 2019, 14, 883–890.

(27) Ehlerding, E. B.; Chen, F.; Cai, W. Biodegradable and Renal Clearable Inorganic Nanoparticles. Adv. Sci. 2016, 3, 1500223.

(28) Boal, A. K.; Ilhan, F.; DeRouchey, J. E.; Thurn-Albrecht, T.; Russell, T. P.; Rotello, V. M. Self-Assembly of Nanoparticles into Structured Spherical and Network Aggregates. Nature 2000, 404, 746–748.

(29) Al Zaki, A.; Joh, D.; Cheng, Z.; De Barros, A. L. B.; Kao, G.; Dorsey, J.; Tsourkas, A. Gold-Loaded Polymeric Micelles for Computed Tomography-Guided Radiation Therapy Treatment and Radiosensitization. ACS Nano 2014, 8, 104–112.

(30) M. Higbee-Dempsey E.; Amirshaghaghi, A.; J. Case M.; Bouché, M.; Kim, J.; P. Cormode D.; Tsourkas, A. Biodegradable Gold Nanoclusters with Improved Excretion Due to pH-Triggered Hydrophobic-to-Hydrophilic Transition. J. Am. Chem. Soc. 2020, 142, 7783–7794.

(31) Tam, J. M.; Tam, J. O.; Murthy, A.; Ingram, D. R.; Ma, L. L.; Travis, K.; Johnston, K. P.; Sokolov, K. V. Controlled Assembly of Biodegradable Plasmonic Nanoclusters for Near-Infrared Imaging and Therapeutic Applications. ACS Nano 2010, 4, 2178–2184.

(32) Cheheltani, R.; Ezzibdeh, R. M.; Chhour, P.; Pulaparthi, K.; Kim, J.; Jurcova, M.; Hsu, J. C.; Blundell, C.; Litt, H. I.; Ferrari, V. A.; Allcock, H. R.; Sehgal, C. M.; Cormode, D. P. Tunable, Biodegradable Gold Nanoparticles as Contrast Agents for Computed Tomography and Photoacoustic Imaging. Biomaterials 2016, 102, 87–97.

(33) Rengan, A. K.; Bukhari, A. B.; Pradhan, A.; Malhotra, R.; Banerjee, R.; Srivastava, R.; De, A. In Vivo Analysis of Biodegradable Liposome Gold Nanoparticles as Efficient Agents for Photothermal Therapy of Cancer. Nano Lett. 2015, 15, 842–848.

(34) Cassano, D.; Santi, M.; D’Autilia, F.; Mapanao, A. K.; Luin, S.; Voliani, V. Photothermal Effect by NIR-Responsive Excretable Ultrasmall-in-Nano Architectures. Mater. Horizons 2019, 6, 531–537.

(35) Zhou, M.; Du, X.; Wang, H.; Jin, R. The Critical Number of Gold Atoms for a Metallic State Nanocluster: Resolving a Decades-Long Question. ACS Nano 2021, 15, 13980–

(36) Stover, R. J.; Joshi, P.; Yoon, S. J.; Murthy, A. K.; Emelianov, S.; Johnston, K. P.; Sokolov, K. V. Biodegradable Plasmonic Nanoparticles: Overcoming Clinical Translation Barriers. In Optical Molecular Probes, Imaging and Drug Delivery, OMP 2015; 2015.

(37) Zaki, A. Al Hui, J. Z.; Higbee, E.; Tsourkas, A. Biodistribution, Clearance, and Toxicology of Polymeric Micelles Loaded with 0.9 or 5 nm Gold Nanoparticles. J. Biomed. Nanotechnol. 2015, 11, 1836–1846.

(38) Azubel, M.; Kornberg, R. D. Synthesis of Water-Soluble, Thiolate-Protected Gold Nanoparticles Uniform in Size. Nano Lett. 2016, 16, 3348–3351.

(39) Knittel, L. L.; Schuck, P.; Ackerson, C. J.; Sousa, A. A. Zwitterionic Glutathione Monoethyl Ester as a New Capping Ligand for Ultrasmall Gold Nanoparticles. RSC Adv. 2016, 6, 46350–46355.

(40) Sánchez-Iglesias, A.; Grzelczak, M.; Altantzis, T.; Goris, B.; Pérez-Juste, J.; Bals, S.; Van Tendeloo, G.; Donaldson, S. H.; Chmelka, B. F.; Israelachvili, J. N.; Liz-Marzán, L. M. Hydrophobic Interactions Modulate Self-Assembly of Nanoparticles. ACS Nano 2012, 6, 11059–11065.

(41) Huang, P.; Lin, J.; Li, W.; Rong, P.; Wang, Z.; Wang, S.; Wang, X.; Sun, X.; Aronova, M.; Niu, G.; Leapman, R. D.; Nie, Z.; Chen, X. Biodegradable Gold Nanovesicles with an Ultrastrong Plasmonic Coupling Effect for Photoacoustic Imaging and Photothermal Therapy. Angew. Chemie Int. Ed. 2013, 52, 13958–13964.

(42) Kwon, N.; Oh, H.; Kim, R.; Sinha, A.; Kim, J.; Shin, J.; Chon, J. W. M.; Lim, B. Direct Chemical Synthesis of Plasmonic Black Colloidal Gold Superparticles with Broadband Absorption Properties. Nano Lett. 2018, 18, 5927–5932.

(43) Chen, J.; Gong, M.; Fan, Y.; Feng, J.; Han, L.; Xin, H. L.; Cao, M.; Zhang, Q.; Zhang, D.; Lei, D.; Yin, Y. Collective Plasmon Coupling in Gold Nanoparticle Clusters for Highly Efficient Photothermal Therapy. ACS Nano 2022, 16, 910–920.

(44) Mir-Simon, B.; Reche-Perez, I.; Guerrini, L.; Pazos-Perez, N.; Alvarez-Puebla, R. A.; Domingo, M. Universal One-Pot and Scalable Synthesis of SERS Encoded Nanoparticles. Chem. Mater 2015, 27, 950–958.

(45) Jain, P. K.; Huang, W.; El-Sayed, M. A. On the Universal Scaling Behavior of the Distance Decay of Plasmon Coupling in Metal Nanoparticle Pairs: A Plasmon Ruler Equation. Nano Lett. 2007, 7, 2080–2088.

(46) Wustholz, K. L.; Henry, A.-I.; McMahon, J. M.; Freeman, R. G.; Valley, N.; Piotti, M. E.; Natan, M. J.; Schatz, G. C.; Van Duyne, R. P. Structure−Activity Relationships in Gold Nanoparticle Dimers and Trimers for Surface-Enhanced Raman Spectroscopy. J. Am. Chem. Soc. 2010, 132, 10903–10910.

(47) Vujačić, A.; Vasić, V.; Dramićanin, M.; Sovilj, S. P.; Bibić, N.; Milonjić, S.; Vodnik, V. Fluorescence Quenching of 5,5′-Disulfopropyl-3,3′-Dichlorothiacyanine Dye Adsorbed on Gold Nanoparticles. J. Phys. Chem. C 2013, 117, 6567–6577.

(48) Walters, C. M.; Pao, C.; Gagnon, B. P.; Zamecnik, C. R.; Walker, G. C. Bright Surface-Enhanced Raman Scattering with Fluorescence Quenching from Silica Encapsulated J-Aggregate Coated Gold Nanoparticles. Adv. Mater. 2018, 30, 1705381.

(49) Silva, W. R.; Keller, E. L.; Frontiera, R. R. Determination of Resonance Raman Cross-Sections for Use in Biological SERS Sensing with Femtosecond Stimulated Raman Spectroscopy. Anal. Chem. 2014, 86, 7782–7787.

(50) Blackie, E. J.; Le Ru, E. C.; Etchegoin, P. G. Single-Molecule Surface-Enhanced Raman Spectroscopy of Nonresonant Molecules. J. Am. Chem. Soc. 2009, 131, 14466–14472.

(51) Nam, J.; Won, N.; Jin, H.; Chung, H.; Kim, S. pH-Induced Aggregation of Gold Nanoparticles for Photothermal Cancer Therapy. J. Am. Chem. Soc. 2009, 131, 13639–13645.

(52) Hou, H.; Zhao, Y.; Li, C.; Wang, M.; Xu, X.; Jin, Y. Single-Cell pH Imaging and Detection for PH Profiling and Label-Free Rapid Identification of Cancer-Cells. Sci. Rep. 2017, 7, 1759.

(53) Song, J.; Kim, J.; Hwang, S.; Jeon, M.; Jeong, S.; Kim, C.; Kim, S. “‘Smart’” Gold Nanoparticles for Photoacoustic Imaging: An Imaging Contrast Agent Responsive to the Cancer Microenvironment and Signal Amplification via pH-Induced Aggregation. Chem. Commun 2016, 52, 8287.

(54) Poon, W.; Zhang, Y.-N.; Ouyang, B.; Kingston, B. R.; Wu, J. L. Y.; Wilhelm, S.; Chan, W. C. W. Elimination Pathways of Nanoparticles. ACS Nano 2019, 13, 5785–5798.

(55) Balfourier, A.; Luciani, N.; Wang, G.; Lelong, G.; Ersen, O.; Khelfa, A.; Alloyeau, D.; Gazeau, F.; Carn, F. Unexpected Intracellular Biodegradation and Recrystallization of Gold Nanoparticles. Proc. Natl. Acad. Sci. 2020, 117, 103–113.

(56) Zavaleta, C. L.; Garai, E.; Liu, J. T. C.; Sensarn, S.; Mandella, M. J.; Van de Sompel, D.; Friedland, S.; Van Dam, J.; Contag, C. H.; Gambhir, S. S. A Raman-Based Endoscopic Strategy for Multiplexed Molecular Imaging. Proc. Natl. Acad. Sci. 2013, 110, E2288–E2297.

(57) Harmsen, S.; Rogalla, S.; Huang, R.; Spaliviero, M.; Neuschmelting, V.; Hayakawa, Y.; Lee, Y.; Tailor, Y.; Toledo-Crow, R.; Kang, J. W.; Samii, J. M.; Karabeber, H.; Davis, R. M.; White, J. R.; van de Rijn, M.; Gambhir, S. S.; Contag, C. H.; Wang, T. C.; Kircher, M. F. Detection of Premalignant Gastrointestinal Lesions Using Surface-Enhanced Resonance Raman Scattering–Nanoparticle Endoscopy. ACS Nano 2019, 13, 1354–1364.

